# Human RNase 2 is essential for macrophage response to viral RNA

**DOI:** 10.64898/2025.12.03.691791

**Authors:** Jiarui Li, Lu Lu, Raúl Anguita, Esther Julián, Ester Boix

**Author notes:** Corresponding authors: Ester Boix,; Jiarui Li.

## Abstract

RNase 2 is the most abundant human RNase A member in macrophages and its expression is activated upon exposure to viruses. Here, we explored the protein role by co-transcriptomics analysis of wild-type (WT) and RNase 2-knock-out (KO) macrophages in absence/presence of a virus-derived single-stranded RNA (ssRNA40). Results revealed that RNase 2 is key for maintaining cell homeostasis. Lacking RNase 2 induced the expression of stress-response markers under basal conditions and abolished the antiviral response of cells exposed to ssRNA40. In contrast, the up-regulated genes in WT macrophages participate in pro-inflammatory signaling response through TLR8-dependent pathways and antiviral immunity, with activation of MAPK and JAK-STAT pathways. Complementarily, we identified five top tRNA-derived small RNAs (tDRs) in response to ssRNA40 related to RNase 2, showing a preferential cleavage sites at CA and uridine rich regions of anticodon loops. Results highlight the essential roles of RNase 2 in antiviral response and inflammatory processes.

## INTRODUCTION

Human RNases from the RNase A superfamily are innate immunity proteins activated upon diverse cellular stress conditions, such as infection. Within this family, antimicrobial RNases constitute a group of small secretory proteins expressed by innate cells that participate in body fluids protection against pathogens.^1–3^ Their secretion to the extracellular space is selectively induced based on the type of external stimuli. Interestingly, whereas bacterial infection specifically induces protein secretion of some family members, such as RNases 3, 6 and 7,^4,5^ RNase 2 is selectively expressed upon viral infection.^6,7^ RNase 2 can inhibit the replication of ssRNA viruses, such as respiratory syncytial virus (RSV)^8,9^ and human immunodeficiency virus (HIV)-1.^6,10^ RNase A family members also differ in their secretory cell type. Human RNase 2, originally called eosinophil derived neurotoxin (EDN) upon its discovery,^11,12^ is a protein mostly expressed in bone marrow and lymphoid tissues (<u>The Human Protein Atlas</u>). The protein expression profile correlated to innate immune response of myeloid leukemia cell lines^13,14^ and comparative cell line profiling (https://celldive.dsmz.de/rna/ll-100) highlighted THP-1 line as abundantly expressing RNase 2, with non-significant or reduced expression of any other RNase A family members.

In our previous work we reported how THP-1 cells-derived macrophages infected by RSV activated RNase 2 expression and how the protein gene knock-out impaired significantly the cell survival to infection.^15^ RNase 2 is also upregulated in SARS-CoV infection^16^ and the protein levels were proposed as a biomarker for COVID-19. Unfortunately, RNase 2 presence was also linked to inflammation at the respiratory tract and severity of clinical symptoms.^17^ Very recently, RNase 2 overexpression was associated with hyperinflammation and correlated to the rare pathogenesis risk following mRNA vaccination.^18^ The authors suggested that cleavage of foreign RNA by RNase 2 released oligonucleotide products bearing uridine 2’3’ cyclic (U>p) ends, which would in turn activate TLR8 receptor, in accordance with previous studies that correlated RNase catalytic activity on foreign RNA with the release of U>p end and endosomal TLR activation to trigger host response.^19,20^ Thus, the RNase protein might have a dual role, working as an alarmin during acute infection and contributing to tissue infection clearance; whereas severe tissue damage and life-threatening hyperinflammation can occur in chronic disease conditions.^1^

In this work, we were interested in understanding how RNase 2 activity mediates macrophage response when cells are exposed to viral RNA. Towards this goal, we performed an integrative analysis of coding and non-coding transcriptome populations in wild-type (WT) and RNase 2 knock-out (KO) THP-1 cells in the absence and presence of the single stranded RNA40 (ssRNA40), a 20mer taken from the HIV-1 genome. Last, to identify specific ncRNAs associated to RNase 2 mediated antiviral response, we performed a selective 2’3’ C>p amplification of small RNA products generated by an endonuclease cleavage.

## RESULTS

### RNase 2 plays a crucial role in maintaining macrophage homeostasis

To explore the role of RNase 2 in macrophage, we performed a transcriptomics analysis of WT THP-1 cells-derived macrophages (R2WT) *versus* the edited RNase 2-KO cell line (R2KO). From the principal component analysis (PCA), we confirmed that R2KO cells expressed genes significantly differentiate from the R2WT ones (Figures S1A and S1B). The overall KEGG (<u>Kyoto Encyclopedia of Genes and</u> <u>Genomes</u>) pathway enrichment (differentially expressed genes (DEGs) cut-off: padjusted (padj) < 10^−3^ and ≥ 5 × |Fold Change (FC)|) underlined a central signaling pathway associated to chemokines/cytokines related to cell response to stress conditions (Figures S1C, S1D and S1E).

Following, a second-NGS analysis was performed to confirm the initial identified differences between R2WT and R2KO macrophages. The compiled analysis of overlapped results from both studies (Additional file 1) highlighted the most significant DEGs related to the absence of RNase 2 in macrophages (Figure 1A). Surprisingly, in R2KO macrophages under basal conditions, we observed an enhanced up-regulation (≥ 5 × FC) of genes related to cytokines and chemokines (Figures 1B, 1C and 1D): cytokine receptors in C-X-C motif chemokine ligand (CXCL) family, CC-type chemokines (CCL2, CCL7, CCL8) and Interleukins (IL7, IL33). The observed expression profile is also characteristic of M1-like macrophages, confirmed by upregulation of CD80 and CD86 M1 markers (Figure 1D).

**Figure 1.**
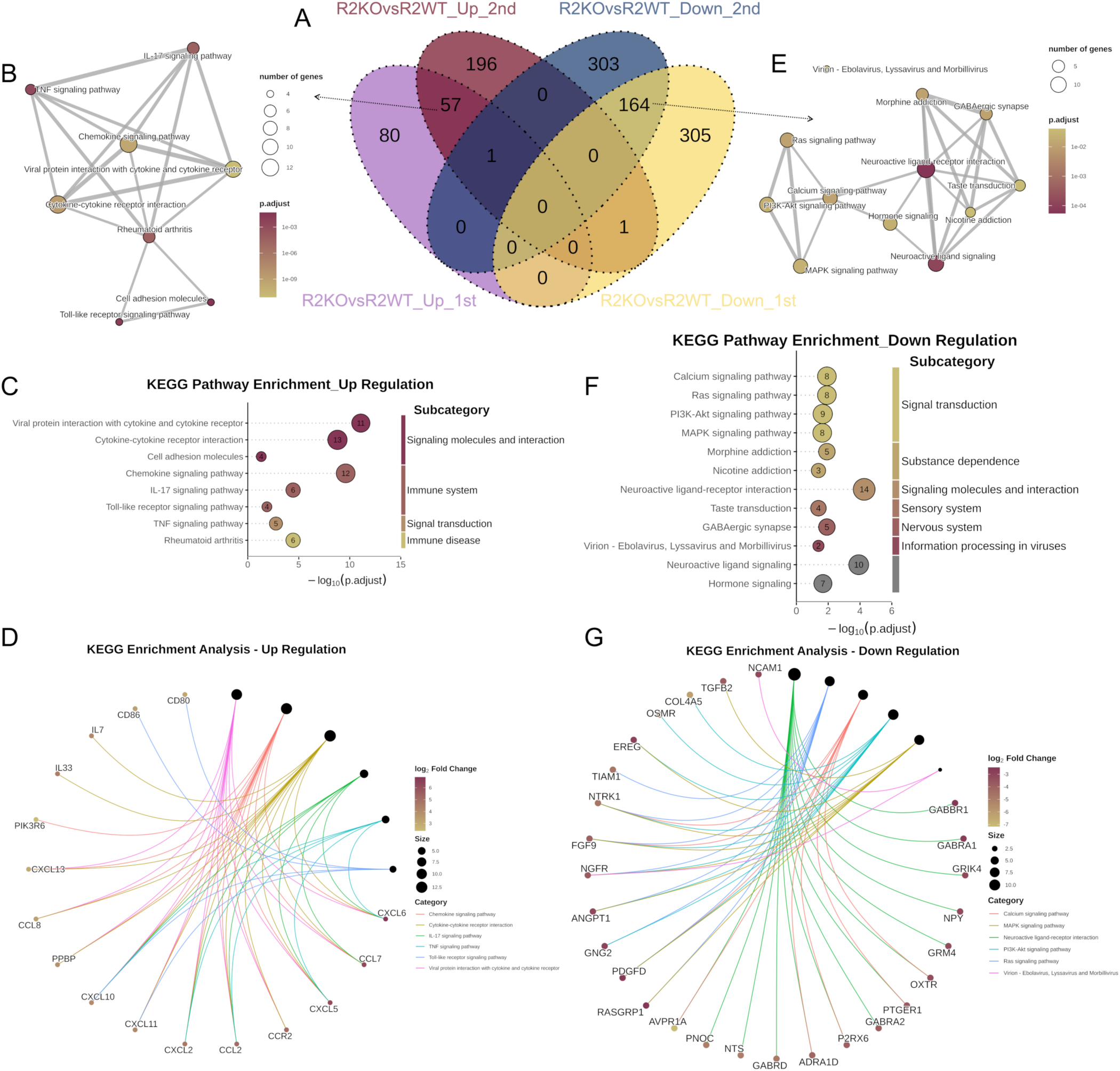
Combined analysis results for 1st and 2nd NGS of R2KO and R2WT THP-1-derived macrophages. **(A)** Venn plot for R2KO vs R2WT genes of 1st and 2nd NGS. **(B, C and D)** KEGG pathway enrichment emapplot **(B)**, dotplot **(C)** and cnetplot **(D)** of overlapping up-regulated genes. **(E, F and G)** KEGG pathway enrichment emapplot **(E)**, dotplot **(F)** and cnetplot **(G)** of overlapping down-regulated genes. The significantly overlapping up/down-regulated genes (|Log_2_FC|>2.32 and padj<0.01) for R2KO vs R2WT were selected and the significantly KEGG pathways (qvalue<0.05) are shown.

On the other hand, the absence of RNase 2 was associated with a significant downregulation (≥ 5 × FC) of genes related to cell cycle regulation pathways. Basically, R2KO cells are significantly deficient in PI3K-Akt, Ras and MAPK markers (Figures 1E, 1F and 1G). Activation of the PI3K/Akt pathway is reported to be critical in restricting proinflammatory and promoting anti-inflammatory responses in TLR-stimulated macrophages.^21,22^ To note, some transcriptome characteristic genes associated to RNase 2 (R2WT) (Figure 1G), such as *TGFB2* (Transforming Growth Factor beta 2, TGFβ2), *NPY* (Neuropeptide Y)^23^ and *OSMR* (Onconstatin M receptor),^24^ are indicative of a M2-like macrophage, involved in cell survival and proliferation. In contrast, many upregulated genes in R2KO are downstream factors in cellular inflammatory pathways (Figure 1D).

All the findings here indicated that RNase 2 plays a key role in maintaining the macrophage homeostasis and may assist to prevent the cell differentiation to a pro-inflammatory state.

### Integrative analysis of mRNA-miRNAs corroborates the key role of RNase 2 in macrophage homeostasis

To characterize the differential behavior of native and knock-out cell lines, we then analyzed the microRNA (miRNA) population. By comparison of R2KO vs R2WT, we identified in the R2KO group 67 miRNAs significantly changed (padj < 0.05, Additional file 2), among which 13 miRNAs had over 4-time underrepresentation (log_2_FC < −2) whereas only four miRNAs were significantly overrepresented (log_2_FC > 2) in R2KO cells (Figure 2A).

**Figure 2.**
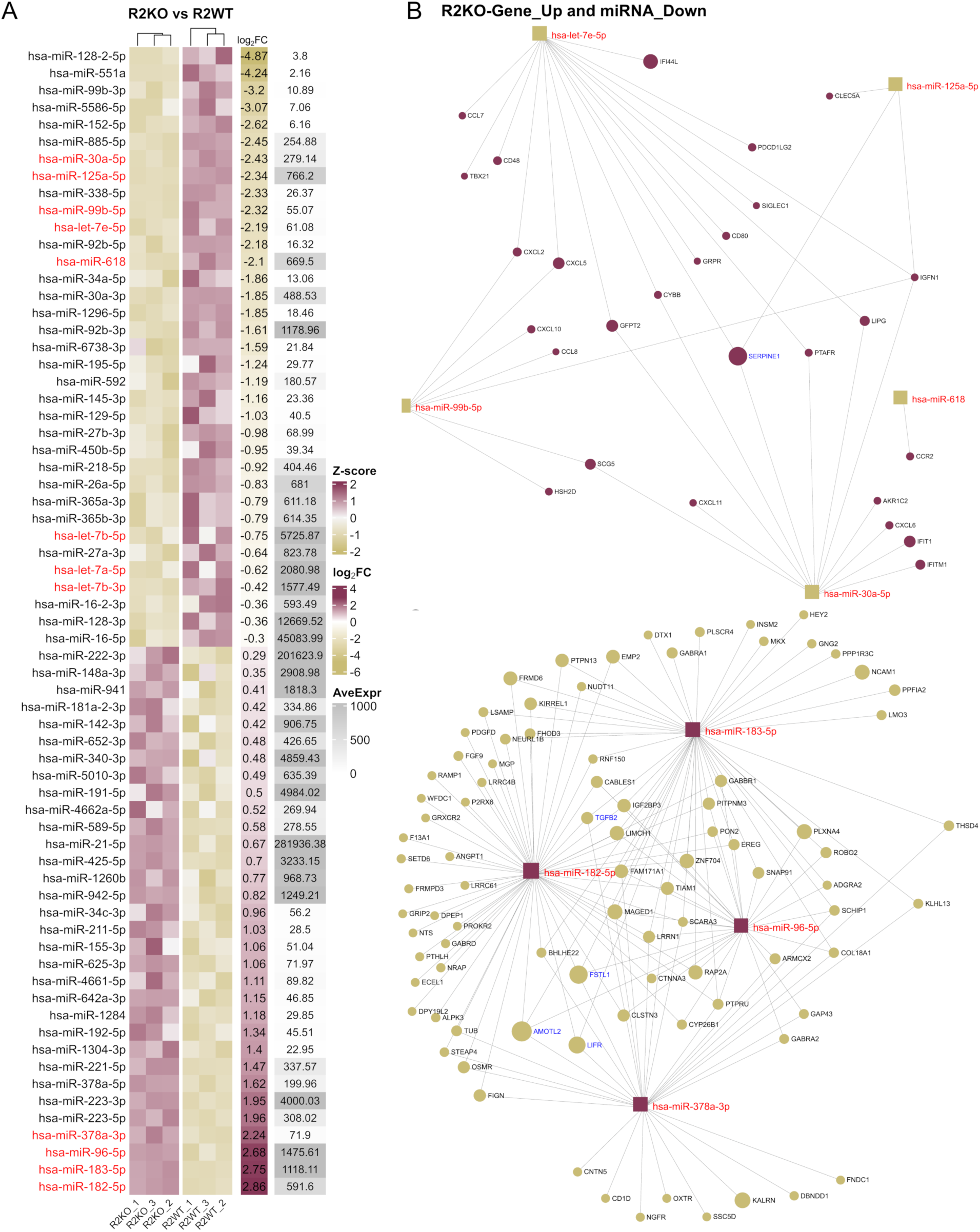
mRNA-miRNA co-analysis of R2KO and R2WT THP-1-derived macrophages. **(A)** Heatmap plot of miRNAs for R2KO vs R2WT. Only over-represented (Up) or under-represented (Down) miRNAs with padj<0.05 were selected. AveExpr: average of normalized counts. **(B and C)** Network analysis of up-regulated genes with under-represented miRNAs **(B)** and down-regulated genes with over-represented miRNAs **(C)** for R2KO vs R2WT comparison. The genes used were the overlapping ones (Figure 1A) from 1st and 2nd NGS. The network was generated through <u>miRNet 2.0</u> (access date: 09/09/2025) by uploading the official gene symbol (|Log_2_FC|>2.32) and miRBase ID (|Log_2_FC| > 2) with Degree filter at 5.0.^29^ Hub miRNA are labelled in red and genes in blue.

From the significant down-representation miRNA members in R2KO vs R2WT, we firstly noticed the let-7 miRNA family, such as hsa-let-7a-5p, hsa-let-7b-5p/3p, and hsa-let-7e-5p. This miRNA family is not only involved in macrophage polarization^25^ but also directly targets viral RNA to inhibit virus replication.^26^ After co-analysis of the most up-regulated genes (R2KO_Gene_Up) (Figure 1A) and down-represented miRNAs (R2KO_miRNA_Down) (Figure 2A), we identified five-top down-represented miRNAs that present a global overlapping interaction with genes most upregulated in R2KO cells (Figure 2B). Among them, hsa-miR-99b, let-7e and miR-125a belong to a cluster associated with the Map kinase p42/44 and p38.^27^ The inhibition of this miRNA cluster resulted in a p42/44 kinase reduction and a p38 activation, thereby disrupting the balance of cytokines production. This result is also consistent with what we observe in the R2KO macrophages, showing down-representation of both the miRNA cluster (Figure 2A) and MAPK signaling pathway (Figure 1G). Apart from miRNAs correlating with genes involved in pro-inflammatory pathways, we also observed a central hub gene (*SERPINE1*) related to regulation of blood coagulation, SARS-CoV-2 signaling and COVID19 adverse outcome (Figures 2B and S2), as identified by WikiPathways analysis.^28^ Results indicate that in R2WT, the identified miRNAs are up-represented and could target the mRNA for Serpine1, blocking its translation. *SERPINE1* codes for a serine protease inhibitor that regulates blood clotting and ensures the proper hemostasis. Thus, the present data suggests that RNase 2 is involved in a proper blood clotting regulation.

Conversely, there were only 4 over-represented miRNAs (4 × FC) in R2KO macrophages, and the microRNA-183/96/182 cluster was significantly identified. The network analysis with the 164 down-regulated genes in R2KO (Figure 1A) also showed well targeted-connection of these 4 miRNAs (Figure 2C) and we found central hub genes including *AMOTL2* (Angiomotin Like 2), *FSTL1* (Follistatin-like 1), *LIFR* (Leukemia Inhibitory Factor Receptor) and *TGFB2*. All of them are related to cell proliferation, differentiation and survival (<u>The Human Protein Atlas</u>).

We further performed an integrated KEGG pathways analysis based on the up-regulated genes and all significant underrepresented miRNAs in R2KO macrophages (Figure 2A). The alluvial plot (Figure 3A) clearly highlighted that these miRNAs target not only *CD80*, *CD86* but also *CCL7* and *IL7*. Consistently, the most significant KEGG pathways from miRNA and mRNA overlapping confirmed the mRNA transcriptomic analysis, where the upregulation of chemokine signaling pathway and cytokine-cytokine receptor interaction were observed (Figure 1D). Thus, we can corroborate an overall pro-inflammatory response profile in the KO cell line.

**Figure 3.**
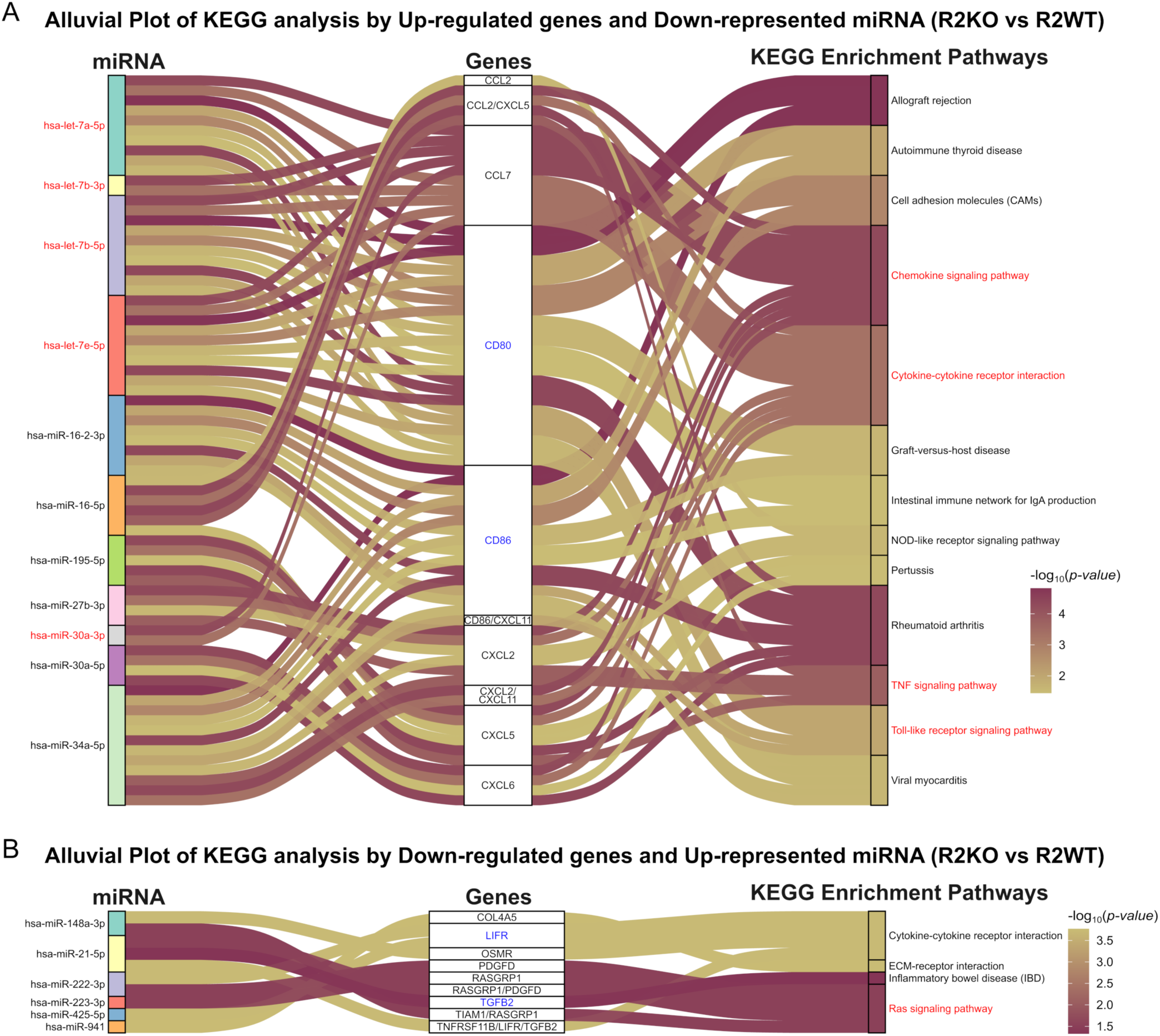
Genes-miRNAs incorporated in KEGG pathways. **(A and B)** The overlapping genes from 1st and 2nd NGS and miRNAs which are up-regulated genes with underrepresented miRNAs **(A)** and down-regulated genes with overrepresented miRNAs **(B)** by R2KO vs R2WT comparison were uploaded into <u>mirPath v.3</u>^30^ (access date: 09/09/2025) to perform the KEGG analysis by TarBase v7.0 with P-value threshold: 0.05 and MicroT threshold: 0.8.

In contrast, we also identified significant over-represented miRNAs (Figure 2A) associated to 164 down-regulated genes (Figure 3B), further confirming that the upregulated miRNAs in R2KO would interfere in transcription levels of genes such as *TGFB2* and *LIFR*, which are important M2-macrophage regulators, as demonstrated in previous DEGs analysis (Figure 1G).

### RNase 2 mediates an antiviral and pro-inflammatory response of macrophages in presence of ssRNA40

Next, we wanted to explore the RNase 2 role during host response to viral exposure. In previous section, we demonstrated cellular metabolic dysregulation after knocking out RNase 2 in THP-1-derived macrophages. However, these data were limited to basal conditions. Interestingly, comparison of RNase 2 WT and KO also highlighted genes related to antiviral response. To corroborate RNase 2 essential role in macrophage antiviral response, we performed a cell exposure assay using ssRNA40 derived from the HIV-1 genome.^31^ Previous works validated the efficacy of the RNA40 oligonucleotide to stimulate macrophage antiviral response.^19,20,32^ Following, we focused on the comparison between both cell lines, R2WT and R2KO in the presence of 4 μg/mL ssRNA40, named WT40 and KO40, respectively.

The PCA analysis of all four transcriptomes confirmed the good discrimination among all groups, with a maximum differentiation between R2WT and WT40 and nearly no significant differences between R2KO and KO40 (Figures 4A-C and S3A-F), indicating macrophages lacking RNase 2 are mostly deprived of the ability to respond to viral RNA exposure.

**Figure 4.**
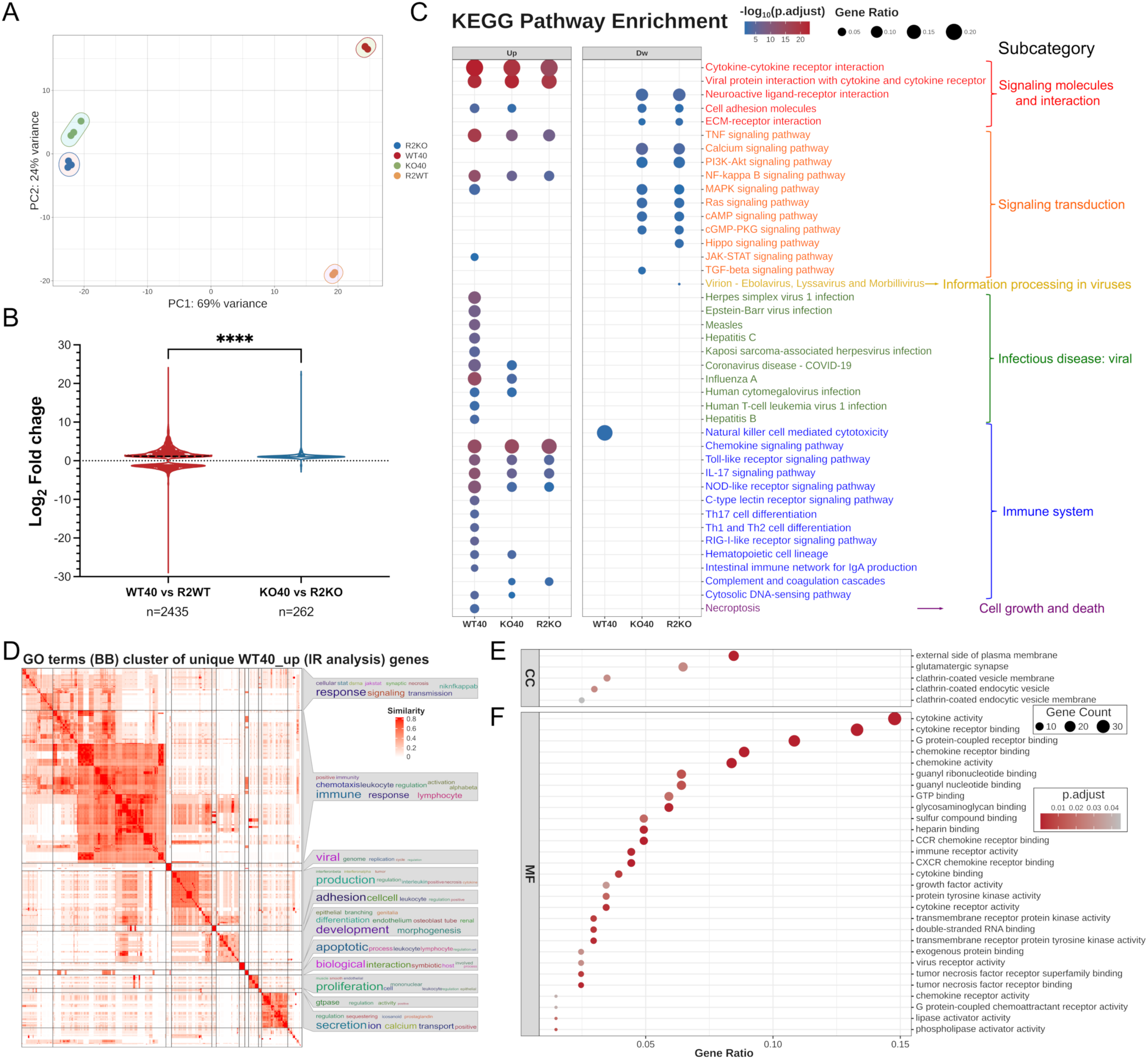
Results of NGS for R2WT and R2KO THP-1-derived macrophage exposed to ssRNA40. **(A)** PCA plot for R2WT and R2KO macrophage cells and R2WT/R2KO exposed to ssRNA40 (WT40/KO40, respectively) of mRNA sequencing. **(B)** Comparison of the distribution of differentially expressed transcripts (Cut-off: (|Log_2_FC|>1 and padj<0.01) between WT40 vs R2WT and KO40 vs R2KO. Kolmogorov-Smirnov test was used to do the statistical analysis. *\*\*\*\*p* < 0.0001. **(C)** Dotplot of KEGG pathway enrichment cluster (qvalue<0.05) of protein-coding genes for WT40, KO40 or R2KO *versus* R2WT. The genes with cut-off:|Log_2_FC|>2.32 and padj<0.01) were selected. **(D, E and F)** Go (Gene Ontology) analysis for unique WT40 up-expressed protein-coding genes (IR analysis). Topgo plot **(D)** of Biological Process (BP) analysis and dotplot of Cellular Component (CC) **(E)** and molecular function (MF) **(F)** analysis. The genes with cut-off:|Log_2_FC|>2.32 and padj<0.01) were selected.

To further identify which pathways are involved in the response of ssRNA40-exposed macrophages related to RNase 2, we extracted the most significant changed genes only associated to RNase 2 and ssRNA40 exposure by interaction analysis (IR analysis) through DESeq2 by ∼genotype + condition + genotype:condition model,^33^ where the unique genes related to genotype: R2WT and condition: ssRNA40 were filtered out. There is a major significant group (214 protein-coding genes, 5 × FC) representative of upregulated genes in WT40 (Additional file 3).

A closer inspection of these most significantly upregulated genes (5 × FC) only related to RNase 2 and ssRNA exposure by the cluster analysis of Biological Process (BP) also highlighted that the differential expression genes in the presence of ssRNA40 were most related to immune response and virus life cycle, the similarity of which was almost at 1.0 (Figure 4D). Besides, the Cell Component (CC) analysis (Figure 4E) indicated that differential genes were mostly related to action at the external side of plasma membrane or clathrin-coated vesicles and could be involved in transport and signal transduction in the cell membrane, especially regulating molecule exchange and signal communication between cells and environment through endocytosis pathways. This finding corroborates the response to ssRNA40 in WT40 cells. RNase 2 is classified as an extracellular/endolysosomal protein and the commercial ssRNA40, which is provided with LyoVec™, a cationic lipid-based transfection reagent, can enter cells through the endocytosis pathways. Last, Molecular Function (MF) analysis (Figure 4F) also demonstrated the activation of virus-related immune responses, chemokine/cytokine activity and receptor binding.

Next, we performed KEGG analysis with those unique up-regulated genes found in WT40 group (Additional file 3) to obtain the gene profiles related to RNase 2 upon virus exposure (IR analysis). Overall, the activation by virus exposure was confirmed, as 9 out of total 12 KEGG enriched pathways in the <u>Human</u> <u>Diseases</u>subcategory “Infectious disease: viral” were mapped according to our identified genes (Figure 5A). We also observed the up-regulated genes for interferon-induced antiviral proteins: *MX1* (MX dynamin like GTPase 1, MxA) and *MX2* (MX dynamin like GTPase 2, MxB).^34^ In addition, we identified key immune receptors (RIGI, IFNLR1, IL18R1), intermediaries (OAS1-3, BIRC3) as well as kinases (MAPK10, SRC) or oxidase component (NCF1) and transcription factors (IRF7, NFKB2, RELB, STAT1), all of which involved here in upstream signaling pathways.

**Figure 5.**
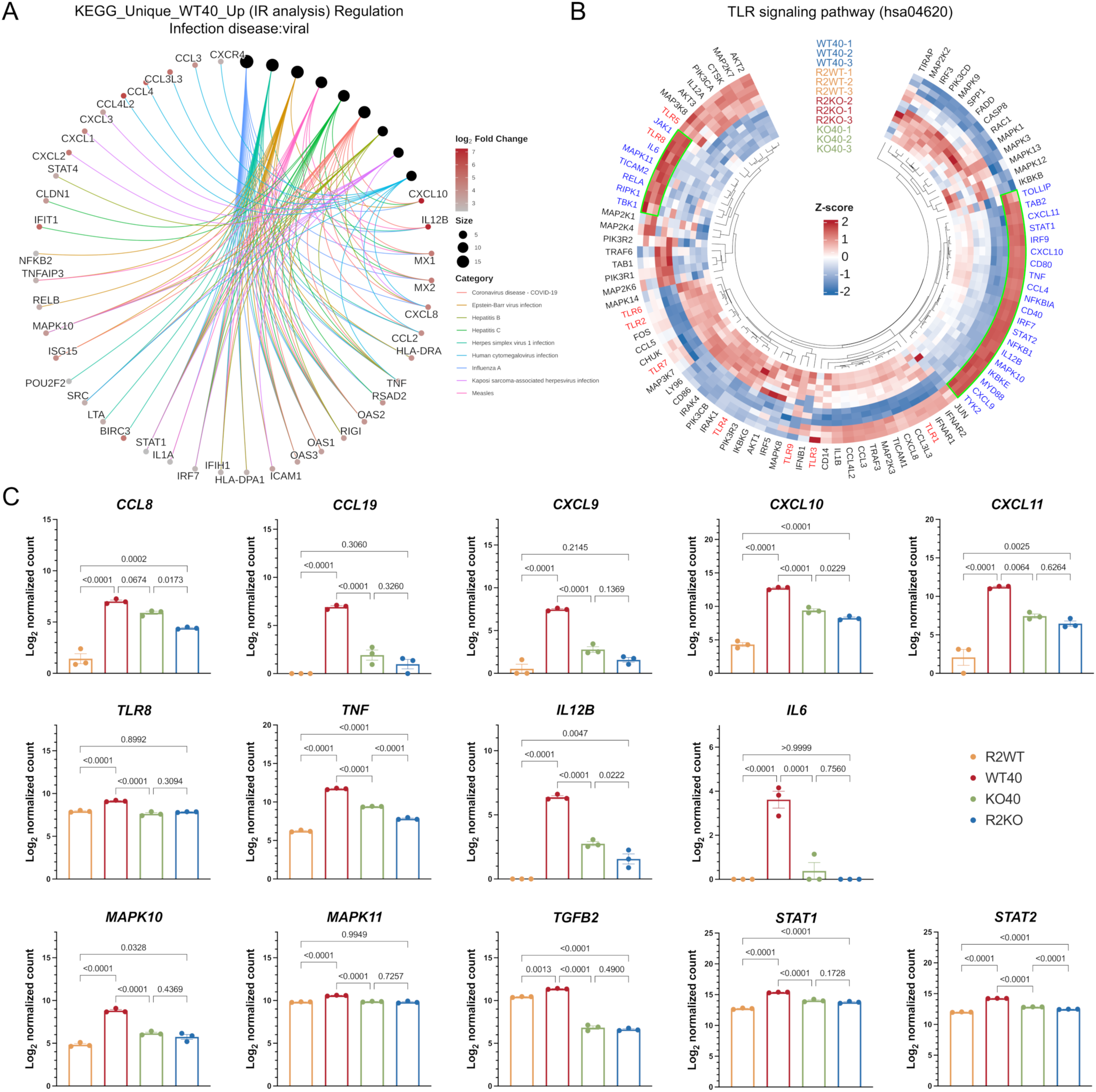
Genes involved in ssRNA40 activation related to RNase 2. **(A)** Cnetplot of KEGG pathways related to viral infections. Only unique WT40 up-regulated genes (IR analysis, log_2_ FC > 2.32 and padj < 0.01) were used. **(B)** Heatmap for genes involved in TLR signaling pathway for all group. The log_2_ normalized count from each sample was used to calculate the Z-score. The different TLRs are highlighted in red and the clusters related to only up-regulated genes in WT40 were highlighted in green squares. **(C)** Log_2_ normalized count of transcripts for represented genes from four groups. Statistical analysis was done by ordinary one-way ANOVA with Tukey-test for multiple comparisons. *p*-value was present between groups.

On the other hand, among the common enrichment pathways that also mapped in the R2KO vs R2WT up-regulation group, such as cytokine-cytokine receptor interaction and Toll-like receptor, TNF or IL-17 signaling pathway (Figures 1D), we identified a specific differentiated profile for the WT40 group. As expected, genes in the CXCL family, CC-type chemokines (Figures 5A, S4A and S4B) not only showed more member diversity but also were upregulated at a much higher level due to activation of ssRNA40 in the presence of RNase 2 (Figure 5C, upper panel). For example, *CCL8* was about 10 times (log_2_FC ≈ 3.28) upregulated in R2KO vs R2WT while over 60 times in WT40 vs R2WT (log_2_FC ≈ 5.96), resulting in an over 20-time change (log_2_FC ≈ 4.42) after applying the IR analysis. Additionally, we found CCL19 and CXCL9-11 chemokines related to adaptive immune response.

Furthermore, we explored how the addition of ssRNA40, a TLR8 RNA agonist, would affect the genes expression in TLR signaling pathway in presence of RNase 2. To this end, the unique genes for WT40 (Addition file 3) involved in TLR-like receptor signaling pathway (hsa04620) were assigned to the pathway view (Figure S5A) and were plotted through the heatmap (Figure 5B) by calculating Z-score across all the group. Results highlight that there are two main clusters of genes that were specifically upregulated in WT40 group. Firstly, it was confirmed that TLR8 was significantly activated in WT40, whereas TLR8 activation in KO40 as well as R2KO were similar to R2WT. Along with the activation of TLR8 signaling pathways, we observed the activation of the NF-κB signaling pathway that leads to the production of antiviral cytokines (Figure S5B). We also found higher activation in WT40 of unique adaptor proteins, MyD88 and TICAM-2 that connect protein to downstream signaling molecules; transcription factors, IRF7, IRF9, REL; and TLR-8-induced cytokines IL-12p70 (*IL12B*), IL-6 and TNF, which are associated to both inflammation and response to virus infection (Figure 5C, middle panel). Together with TLR8 activation, a closest inspection identified key genes involved in kinase pathways: *MAPK10* (JNK3), *MAPK11* (p38β), *TGFB2* (Figure 5C, down panel) participating in MAPK signaling pathway (Figure S5C), which are indicators of macrophage immune response. Finally, we found activation of the JAK-STAT signaling pathway (Figure S5D), with increased expression of associated transcription factors: IRF9, STAT1/STAT2, which mediates IFN antiviral response.

All the results suggest that RNase 2 is essential for macrophage response to viral derived RNA through TLR8 activation and MAPK/JAK-STAT pathways.

### RNase 2 determines the content of tDRs in macrophages in presence of ssRNA40

Non coding-RNAs (ncRNAs) have emerged as key regulators in cell communication and can connect distant cells through vesicle transfer. Vesicular compartment is one of the main identified by CC analysis according to gene enrichment associated to ssRNA40-activated macrophages in presence of RNase 2 (WT40, Figure 4E). We also noticed that miRNA population was significantly altered when knocking-out RNase 2 in macrophages in absence of external stimulus.

Based on the results above, we aimed to further explore how antiviral action by RNase 2 is achieved and whether it is mediated by its direct action on small RNA processing dependent on its ribonucleolytic activity. Before performing a full comparative NGS analysis of ncRNA population on our R2WT, R2KO, WT40 and KO40 cell conditions, we discarded any significant involvement of other cell endoribonucleases. When checking the enrichment of proteins with RNase activity in R2WT vs R2KO, we observed first an over 50-time change (5.69 × Log_2_FC, padj. < 1 × 10^−100^) of RNase 2 but no obvious upregulation (over 10 times) of other RNase A superfamily members or other RNases. Specifically, we observed a similar but slightly higher overexpression of RNase 2 for WT40 vs KO40 (nearly a 60-time change, 5.87 × Log_2_FC, padj. < 1 × 10^−100^) and no significant increase of other RNases, nor of the ubiquitous RNase inhibitor RNH1 (Figure S6). Therefore, we proceeded to explore whether specific ncRNAs processing attributed to WT cell line can be mainly associated to RNase 2 expression in presence of ssRNA40.

To this purpose, apart from polyA mRNA library (Library 1), we analyzed an additional set of three distinct small RNA libraries (Figure S7) that were prepared based on specific amplification procedures: Library 2: Small RNAs (<200nt) with standard 5’P-3’OH ends; Library 3: 2’3’C>p (cP) ends, which are product of an endonuclease cleavage and Library 4: Total small RNAs bearing any 5P/5OH or 3P/3OH/cP ends.^35^ Overall, inspection of these three prepared small RNA libraries highlights that: for Library 2, nearly 40% were derived from long non-coding RNAs (lncRNAs) with about 5% miRNA (Figures S8A-C). The Library 3 was enriched with about 20% tRNAs but only minor miRNAs (< 1%) assigned (Figures S8D-F). For library 4, various small RNAs were identified without extremely high annotated reads for each type (Figures S8G-I). As there was no good discrimination in Library 4 (Figure S8G), and this dataset was merely conceived as an additional control, we pursue our study without further analysis of this data. Therefore, we decided to focus on the other two libraries and specific ncRNA subpopulations: miRNA in Library 2 and tRNA in Library 3, uniquely associated with ssRNA40 activation.

Firstly, an initial comparison across couple groups of miRNA reads in Library 2 was applied (Figures S9A-E). The obvious variance (45%) among the 4 groups was due to the presence or absence of RNase 2 while low variance (15%) was observed upon ssRNA40 exposure (Figure S9F). After overlapping the differently abundant miRNAs by Veen plot, we identified 25 up-represented and 23 down-represented miRNAs related to RNase 2 when macrophages were exposed to ssRNA40 (WT40 vs KO40) that were already present in R2WT vs R2KO (Figure S9G). Although there were 17 unique up-regulated and 9 down-regulated miRNAs in WT40 vs KO40 group (Figure S9H), no significant changes in their relative abundance were observed related to ssRNA40 after IR analysis, suggesting that miRNA may not be the main target of RNase 2 when macrophages are exposed to viral ssRNA.

Next, we explored the RNase 2 potential action on the tRNA population, predominant in Library 3 where the RNase-specific cleaved products (2’3’C>p end) were amplified. In our previous studies,^15,36^ we found that RNase 2 selectively cleaved tRNAs. However, those works focused on the different tRNA-derived small RNAs (tDRs) population between R2WT and R2KO at basal conditions. Here, we wanted to investigate the effect of RNase 2 on tRNA when macrophages are exposed to ssRNA40. Firstly, we analyzed the tRNA isotypes specifically amplified among the four groups in Library 3. tRNA^Glu^ and tRNA^Val^ occupied the two top identified isotypes among the tRNA reads in all groups (around 60-80% totally), together with a slight increase of tRNA^Val^ but decrease of tRNA^Glu^ in R2KO cells (Figure S10A). In addition, the five-prime tDRs were the predominant tRNA fragment type identified in all conditions, with overall values above 60%. In contrast, in RNase 2 knocked out cells in presence of ssRNA40, we observed about a 10% decrease of 5’tRNA fragments and instead, there was a slight increase in other fragment type (Figure S10B). Next, we found some punctual differences in R2KO vs R2WT but the highest significance was observed in the KO40 vs WT40 group (Figures S10C-E). Further analysis of tDRs distribution illustrated that both five-prime and three-prime were the main tDRs in overrepresented WT40 vs KO40 group (Figure S10F) whereas in underrepresented WT40 vs KO40, most different tDRs were defined as “others”, which include all fragments that did not fit to any of the defined categories (whole, five-prime and three-prime). The results indicate that RNase 2 plays a key role in the selective generation of tDRs when macrophages are stimulated by ssRNA40.

After IR analysis that filter out the most distinctive tDRs related to RNase 2 upon ssRNA40 exposure in the macrophages, we identified 17 up-represented and 11 down-represented tDRs (|log_2_ FC|>1, padj<0.05, Additional file 4). Consistent with the two-by-two comparison (Figure S10), most up-represented tDRs in presence of RNase 2 upon ssRNA40 exposure were assigned to five/three-prime. We then mapped the corresponding sequences of these up-represented five/three-prime tDRs to their parental tRNA isotypes according to the unique coverages output from tRAX pipeline in WT40 group. There were 8 dominant tDRs easily tracked by the cleavage site from original sequences (Figure S11). The types of tDRs were also in agreement with the directly statistical analysis by using the unique tDRs count from tRAX output (Figure 6).

**Figure 6.**
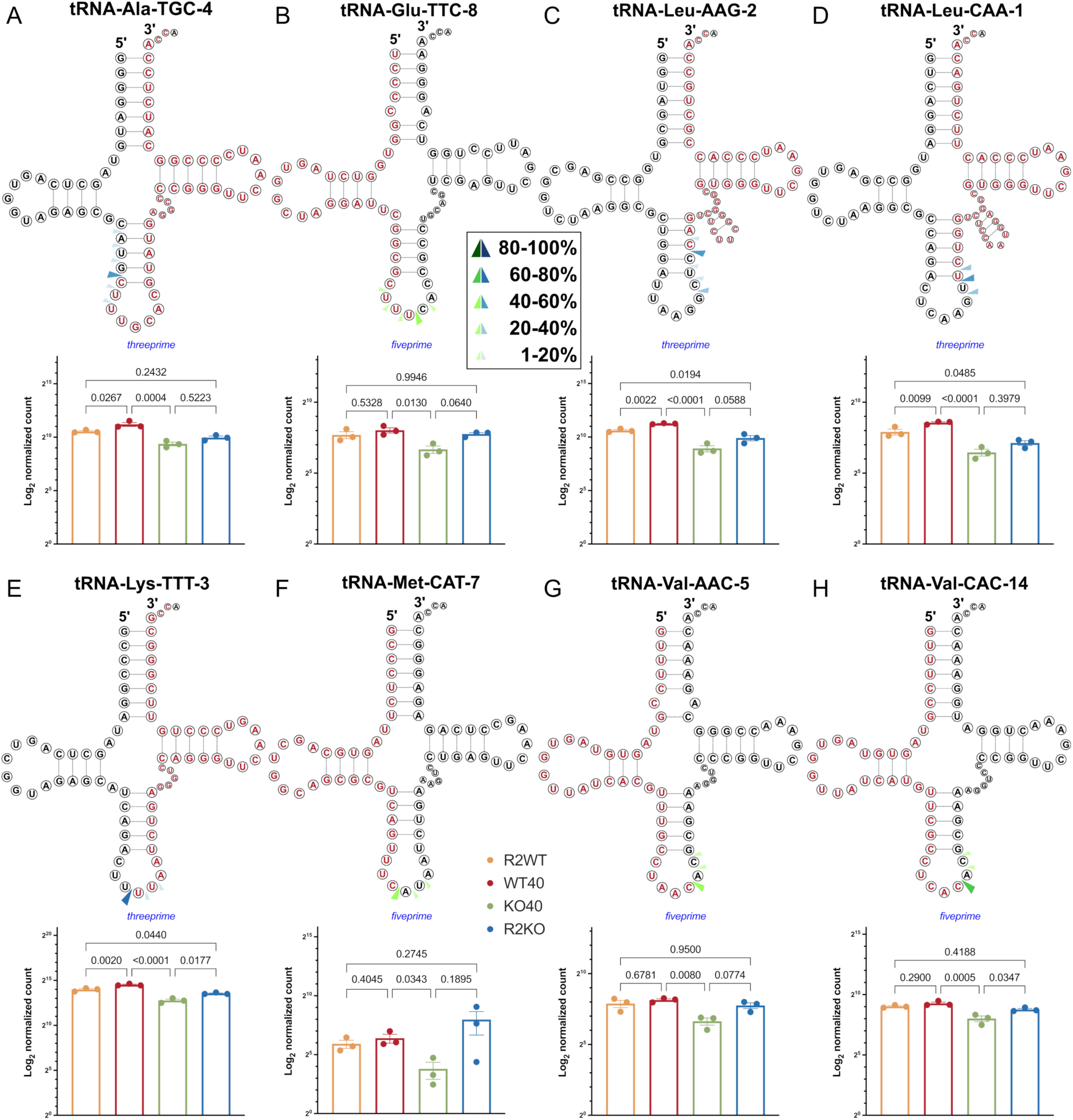
Main identified tDRs related to RNase 2 upon ssRNA40 exposure. Parental tRNAs with significant coverage differences between R2WT and R2KO macrophage cells after 14h ssRNA40 exposure (WT40 and KO40, respectively) are depicted and the identified tDRs are marked in red. The possible cleavage sites were based on the 3′-terminal positions of the five prime fragments (green arrows) and the 5′ terminal positions of three prime fragments (blue arrows) according to the different coverage (Figure S11). The log_2_ normalized read count for all the 4 group were plotted under the corresponding tRNA (tRAX output). Statistical analysis was done by ordinary one-way ANOVA with Tukey-test for multiple comparisons. *p*-value was present between groups. Only sequences with a log_2_FC > 1.0 and padj < 0.05 were included. The detailed of main fragments and their corresponding tDRs types are listed in Table S1.

Firstly, the results showed that 5 out of 8 tRNAs have significant coverage differences at the anticodon loop, resulting in tRNA halves, also known as tRNA-derived stress-induced RNAs (tiRNAs). The coverage of the other three tDRs (see Table S1 for their corresponding names), tDR-31:75-Ala-TGC-4 (Figure 6A), tDR-41:75-Leu-AAG-2 (Figure 6C) and tDR-39:75-Leu-CAA-1(Figure 6D), showed differences at the stem region close to anticodon loop. Secondly, the 5’ tRNA halves, tDR-1:34-Met-CAT-7 (Figure 6F), tDR-1:36-Val-AAC-5 (Figure 6G) and tDR-1:36-Val-CAC-14 (Figure 6H) revealed cleavage at CA from the parental tRNAs, consistent with our previous study where RNase 2 cleaves tRNAs with base specificity at B1 (U/C) and B2 (A) sites.^36^ Moreover, the tRNA^MetCAT^ was also present in the array analysis of our previous study where R2WT and R2KO cell lines were exposed to the respiratory syncytial virus (RSV).^15^ On the other hand, tDR-1:35-Glu-TTC-3-M3 (Figure 6B), tDR-35:75-Lys-TTT-3 (Figure 6E) and tDR-39:75-Leu-CAA-1 (Figure 6D) have uridine or di/tri-uridines at either 3’terminal or 5’terminal.

Therefore, from Library 3 where 2’3’C>p ends product of an endonuclease cleavage were amplified, we can conclude that RNase 2 determine the content of tDRs and mainly produce specific tRNA halves in macrophages in presence of ssRNA40, with preferential cleavage at CA and uridine rich regions. From the predominant eight tDRs, according to the statistical and coverage analysis, we also highlighted the tDR-35:75-Lys-TTT-3, a top tDRs with the highest significance between WT40 vs KO40 (p < 0.0001) and a unique coverage difference above 1000 in comparison to 100 reads for the other tDRs.

## DISCUSSION

RNase 2 is expressed in diverse blood cell types, and its expression is induced upon virus infection, with a reported broad antiviral activity on ssRNA viruses.^1,6^ Although human RNase 2 biological properties were previously associated to host antiviral response, no characterization was performed on the mechanism of action of the protein expressed by macrophages. Macrophages are key immune cells in host response against viral infections. Immune response to viral infection can also lead to inflammation and the process has been associated with ssRNA-triggered TLR8 activation.^37^ On the other hand, TLR8 activation by foreign RNA degradation products was recently reported,^19,20^ where RNase 2 was specifically referred as one of the major actors at endosomal level involved in pathogen RNA cleavage and macrophage response induction.^20^ However, an exacerbated pro-inflammatory response associated to COVID19 mRNA vaccination was reported, pointing to RNase 2 as the triggering molecule.^18^ RNase 2 was also directly correlated to infective endocarditis and sepsis.^38^ Thus, it is important to elucidate RNase 2 regulation pathways to design novel safe therapies. Here, we have further explored the pathways that mediate RNase 2 action in presence of foreign RNA, using THP-1-derived macrophages exposed to ssRNA40. THP-1 cells are a major source of RNase 2 in basal conditions and the current transcriptomics analysis of THP1-derived macrophages confirmed RNase 2 as the main expressed RNase (Figure S6).^20,39^

Firstly, we compared the THP-1 control cells with a RNase 2-knockout cell line, previously obtained in our laboratory.^15^ Comparison of native RNase 2-WT cells (R2WT) *versus* RNase 2-knock out cells (R2KO) highlighted the PI3K-Akt and MAPK signaling pathways, unique to RNase 2 (Figures 1E and 1F). Interestingly, other RNase A family members were previously associated with the PI3K-Akt pathway, such as human RNase 7 and mice eosinophil associated RNases.^40,41^ Specifically, RNase 7 together with other antimicrobial peptides can activate the PI3K path, contributing to relieve skin inflammatory disorders.^42^ The PI3K-Akt pathway is involved in macrophage type 2 differentiation (M2), which participates in cell anti-inflammatory/anti-infective activities. For example, the pI3K/Akt and MAPK pathways are related to the inhibition of MERS-CoV replication *in vitro*^43^ and also regulate cell survival and tissue remodeling. Here, among DEGs associated with the presence of RNase 2 we found markers such as *TGFB2* (Figure 1G), which is characteristic of polarization to M2 macrophage profile and tissue repair. Interestingly, previous work on RNase 2 closest homologue (RNase 3, sharing a 70% amino acid identity) reported induction of TGFβ1 associated to wound healing.^44^ In contrast, when analyzing the upregulated DEGs in the RNase 2-knock out cell line (R2KO vs R2WT), we observed a prominent activation of chemokine signaling pathway together with the CD86 and CD80 cell markers characteristic of M1 macrophages (Figures 1B-D). In addition, comparative analysis of miRNA population highlighted the down-representation of miRNAs that target proinflammatory genes in the RNase 2 knockout (Figure 1D), confirming a pro-inflammatory profile.

Moreover, the miRNAs underrepresented in R2KO cells interact with SERPINE1, which codes for plasminogen activator inhibitor-1 (PAI-1), a key protease inhibitor that regulates blood coagulation. The hub central position of *SERPINE1* gene in the mRNA-miRNA co-analysis (Figure 2B) suggests that the presence of RNase 2 will be important to ensure a proper blood clotting process. Previous work on RNase 1-KO mice correlated absence of RNase 1 with a higher coagulation rate^45^ compared to WT mice. In that study, the loss of RNase 1 was associated with increased levels of FXIIa and suggested to be due to an increase of free extracellular RNA. In the case of RNase 2, the potential protein involvement in blood clotting regulation requires further examination. Curiously, data analysis at *The Human Biomolecular Atlas Program* (<u>HubMAP</u>) identifies RNase 2 and RNase 3 together with the serine proteases inhibitor *SERPINB10* among the top 10 genes that correlate with asthma and haematopoiesis.^46^ Another study correlates high expression levels of serine protease inhibitors (SERPINs), SERPINA1, SERPINE1 and SERPINE2 to less susceptibility to SARS-CoV-2 infection.^47^ Currently, there is increasing evidence that PAI-1 plays key roles for viral envelope fusion during entry and thereby affects virus spread.^48–50^ Collectively, the loss of *TGFB2-AS1* would enhance the transcription and expression of TGFβ targeting genes, FN1 and SERPINE1 and the authors also pointed that inhibition of p38 and JNK would reduce *TGFB2-AS1* induction.^51^ Here, in this work, either down-regulated *TGFB2* or up-regulated *SERPINE1* are the central hubs clustered by the over or under-represented miRNAs population, and *TGFB2-AS1* is also significantly down-represented in case of macrophages lacking RNase 2. On the other hand, the overrepresented miRNAs in R2KO cells, mainly the microRNA-183/96/182 cluster (Figure 2C), is reported to be a key mediator of innate and adaptive immunity, where dysregulation is associated to immune disorders such as autoimmunity. The upregulation of the microRNA-183/96/182 cluster is associated with inflammation and M1 macrophage polarization.^52^

Overall, activation of cell stress signaling pathways in the R2KO line emphasizes the key role of RNase 2 in maintenance of cell homeostasis. Comprehensive inspection on both up and down regulated genes indicates that lack of RNase 2 induced an abnormal upstream signal and may lead to a cellular immune regulation disorder, which in turn can stimulate the expression of downstream immune response factors. Correlation between RNase activity and inhibition of pro-inflammation is also observed for skin RNase 7 in atopic dermatitis, where RNH1 released by cell damage blocks the anti-inflammatory properties of the protein.^53^

The comparative transcriptome of WT and KO cell lines also correlated the presence of RNase 2 protein to a macrophage characteristic antiviral response. Thus, we decided to confirm whether RNase 2 could directly participate in the cell response to viral RNA. To this purpose, we compared the WT and RNase 2-KO cell lines after exposure to a synthetic ssRNA derived from HIV, which was validated previously as a positive control on TLR sensing.^19,20,54^ Our results demonstrate that R2KO macrophages can hardly respond to viral RNA exposure (Figures 4B and S3F). Next, by integrative transcriptome analysis of WT and R2KO macrophages in presence/absence of ssRNA40, we identified the genes uniquely up-regulated in WT40 group. The immune system activation in the present experimental assay conditions was confirmed by a predominant increase of IL6 (log_2_FC ∼ 4, p = 0,0001) (Figure 5C). Analysis of WT40 KEGG cluster pathways indicates the specific cell differentiation to Th1, Th2 and Th17, all of them characteristic of cell immune response to pathogens and adaptive immunity, uniquely for WT exposed to ssRNA40 (Figure 4C). GO enrichment analysis (Figures 4D-F) also clearly indicated that the unique genes from WT40 were involved in triggering antiviral related signaling pathways in macrophages. Therefore, the present data is the first to discriminate the cell basal condition associated to RNase 2 from its specific cell activation mediated by ssRN40. In addition to initial comparison of R2KO vs R2WT up-regulation group (Figure 1D), where we identified genes characteristic of macrophage M1 polarization, indicators of a cell homeostasis imbalance after RNase 2 deletion, the integrative comparison of all conditions in absence/presence of ssRNA40 revealed the specific contribution of RNase 2 in antiviral response. Interestingly, together with CXCL10, we also observed CCL19 and CXCL9 chemokines significantly up regulated only in WT40 (log_2_FC >5 and p < 0.0001, Figure 5C), indicative of the widespread activation of the adaptive immunity.^55^

Complementarily, a heatmap of genes involved in TLR signaling pathways was performed highlighting a unique specific path associated with TLR8 for the WT40-up group (Figure 5B). We observed how TLR8 clusters with IL-6, an inflammation positive marker, and visualized a main cluster that includes STAT1, STAT2 and IRF9 transcription factors, which are characteristic of antiviral response^56^ and linked to pro-inflammatory genes, such as CXCL10, CXCL11 and TNF also reported for RNase 2^20^ and other secretory RNases, such as RNase 7^57^ and RNase T2.^20^ Other secretory RNases from the RNase A superfamily were also associated with antiviral activity, tissue damage and pathogenesis. To note, RNase 1 levels were reported as good predictors of COVID outcome.^58^ RNase 7, for example, abundantly expressed in skin epithelial cells, is associated with antiviral action against herpes virus pathological conditions,^59^ such as atopic dermatitis.^60^ A very recent study proposed a synergic action between RNase 7 and IL6 that might amplify the host response during skin infection.^61^ Another close homologue to RNase 7, RNase 6, was identified as a hub central gene in the macrophage inflammatory response in diabetic nephropathy pathogenic condition.^62^

Next, by simultaneous analysis of coding and non-coding transcriptome, we explored whether RNase 2 enzymatic activity on small RNA population could mediate the cell response. In our previous work, we characterized the RNase 2 endonuclease cleavage products by comparison of ncRNA population between WT and R2KO THP-1-derived macrophages.^15^ A selective cleavage pattern was identified and corroborated *in vitro* using synthetic tRNAs and designed DNA/RNA hairpins.^36^ On the other hand, comparison of WT-RNase 3 and its catalytic-defective H15A mutant indicated that the enzymatic activity was involved in the protein induction of antiviral response in THP1-derived macrophages.^4^ A previous work using the recombinant protein and site-directed mutagenesis correlated RNase 2 antiviral activity on RSV to its enzymatic action.^9^ Thus, we decided to characterize the generation of specific small RNA products associated with presence of RNase 2 in macrophages.

To unambiguously analyze RNase 2 action in our cell working model, we first discarded contribution of other RNases. We first quantified the expression levels in THP-1 cells of other secretory RNases (Figure S6) from the pancreatic RNase A superfamily together with RNase T2, the other main secretory RNase in macrophages^63^ and RNase L, an intracellular RNase associated to viral response.^64^ Considering the previous knowledge on RNH1-RNase 1:1 binding stoichiometry,^65^ the detected inhibitor expression levels would be sufficient for protection of small RNAs at the cytosol level. To note, the unique genes we identified from polyA transcriptome analysis associated to ssRNA40 activation in presence of RNase 2 (Additional file 4) were related to activity at the external side of plasma membrane or chathrin-coated vesicles (Figure 4E). Thus, results suggested that differences observed in the current sequenced ncRNA population would be mostly ascribed to an RNase activity at the extracellular/ endosomal compartment level.

To explore RNase 2 direct effect on small RNA population, we prepared three small RNA library types by sequence amplification according to their 5’ and 3’ ends. Overall, no major changes in miRNA related to ssRNA40 exposure were observed. In contrast, amplification of small RNAs bearing a 2’3’C>p end, which is associated to an endonuclease cleavage,^66^ highlighted significant changes in tDRs population. Whereas we did not observe a specific enrichment of tRNA types in basal conditions when comparing R2WT vs R2KO cell lines, we did find significant changes upon exposure to ssRNA40. Interestingly, we found a specific set of 8 tDRs, with top predominance among the 2 × FC up-representing ones when macrophages were exposed to ssRNA40 in presence of RNase 2 (Figures 6 and S10). The majority of them were previously related to immune response and/or virus infection. For example, we found Glu-TTC and Val-CAC, previously reported by Das and coworkers associated with cell stress conditions.^67^ The lead tDRs according to discrimination significance and unique coverage is the 3’ tRNA half for Lys^TTT^ (Figures 6 and S11), which was identified in our previous work.^15^ Interestingly, tRNA^LysTTT^ serves as a primer for reverse transcription by HIV viruses.^68^ Recently, significant upregulation of only three tDRs, including the tRNA-Lys^TTT^ derived fragment, have been identified in mice infected by Influenza A virus (IAV).^69^

tDR ends can work as endogenous agonists of TLR7 and TLR8.^19,70^ The present data on selective RNase 2-mediated tRNA cleavage should contribute to identifying the specific tDRs released upon viral RNA exposure. For example, the terminal GUUU sequence of the tRNA^ValAAC/CAC^ 5’ half has been identified as a universal signature for TLR7 activation.^70^ Moreover, the uridine-end tDR-1:35-Glu-TTC-3-M3, tDR-35:75-Lys-TTT-3 and tDR-39:75-Leu-CAA-1 have been suggested as agonists of TLR8.^19^ In our previous comparative work of RNase 2 cleavage specificity,^36^ we highlighted the structural determinants to favor the protein interaction with the tRNA at anticodon loop region, with base selectivity for CA at the main and secondary sites, in accordance with previous work using dinucleotides.^71^ Complementarily, studies using extended single stranded homopolymers reported preference of uridine stretches over cytidine for the called “non-pancreatic RNases” group, where RNase 2 is ascribed, in contrast to “pancreatic RNases” group that includes RNase 1, with marked preference for poly-cytidine substrates over poly-uridine.^72^ Other studies^19,20,73^ have also reported cleavage after uridine bases for RNase 2. Here, by inspection of tRNA cleavage by RNase 2 in response to ssRNA40 (Figure 6), we found three tRNA halves cleaved at CA together with cleavage at uridine rich stretches vicinity. Although previous works have reported stress induced tRNA halves release by human RNase 5/angiogenin,^74,75^ angiogenin role in generation of regulatory tDRs was mostly associated to other diseases, such as neurodegenerative processes and cancer.^76,77^ There is also one study that correlated angiogenin generated tDRs with RSV infection.^78^ In that work, the authors reported the tRNA-GluCTC as potential target of RNase 5 and identified a cleavage site with a distinct specificity. Therefore, it is of outmost importance to properly ascribe the distinct cleavage selectivity for each RNase associated with each physiological condition. In this context, activation of endosomal TLR8 by 3’ uridine ends will be associated to RNase 2 but not to RNase 5/ANG, which is expressed by distinct cell type upon other disease conditions and shows a differentiated substrate preference.^1,79^

The present data highlights the key role of RNase 2 expressed in macrophages and the specific tRNA cleavage upon cell exposure to viral RNA. An exponential growth of studies has pointed out changes of ncRNA population especially tDRs during viral infection, as mostly reported for SARS-CoV-2.^80–82^ Some works recently demonstrated that the significant changes in tDRs population in severe COVID-19 may be attributed to potential action of selective RNases cleavage.^83^ Specific 5’tRNA fragments have been detected at sputum and blood samples, associated with angiogenin activity in asthma pathogenesis leading to propose them as disease markers.^84^ The authors reported the predominance of few specific tRNAs isotypes (tRNA Glu, Gly, Lys and Val), three of which were found among our tRNA fragment list associated to RNase 2 presence (Figure 6). The population of tRNA generated products by diverse cell RNases is emerging as a new cellular information code system that participates in cell defense, adaptation to its environment and cell-to-cell signaling.^1,54,85–87^ Moreover, some tDRs have been proposed as disease biomarkers^88^ and even therapeutic molecules.^89^

In summary, by integrative coding and non-coding transcriptomics we show that deletion of RNase 2 in THP-1-derived macrophages disrupted the cell homeostasis. Besides, we prove the direct protein involvement in activation of the PI3K/Akt pathway. Importantly, our results demonstrate that removal of RNase 2 nearly abolished macrophage response to viral ssRNA. The pathways uniquely upregulated by WT in presence of the ssRNA40 were characteristic of TLR8 activation and the STAT1/STAT2/NFkB pro-inflammatory path. Last, we identified a specific set of five tRNA derived fragments uniquely released upon macrophage stimulation by viral RNA mediated by RNase 2 presence, where the 3’ tDR Lys-TTT stands as the most significant representative. Therefore, based on the above results, we can hypothesize that the generation of specific tDRs reinforce the immune response through TLR 8 activation and the JAK-STAT pathway (Figure 7). The identification of specific RNAs signaling is of particular interest to unravel RNase 2 mechanism of action, which has been found to ensure the macrophage homeostasis and mediate the cell antiviral response, but also contribute to pro-inflammatory conditions.

**Figure 7.**
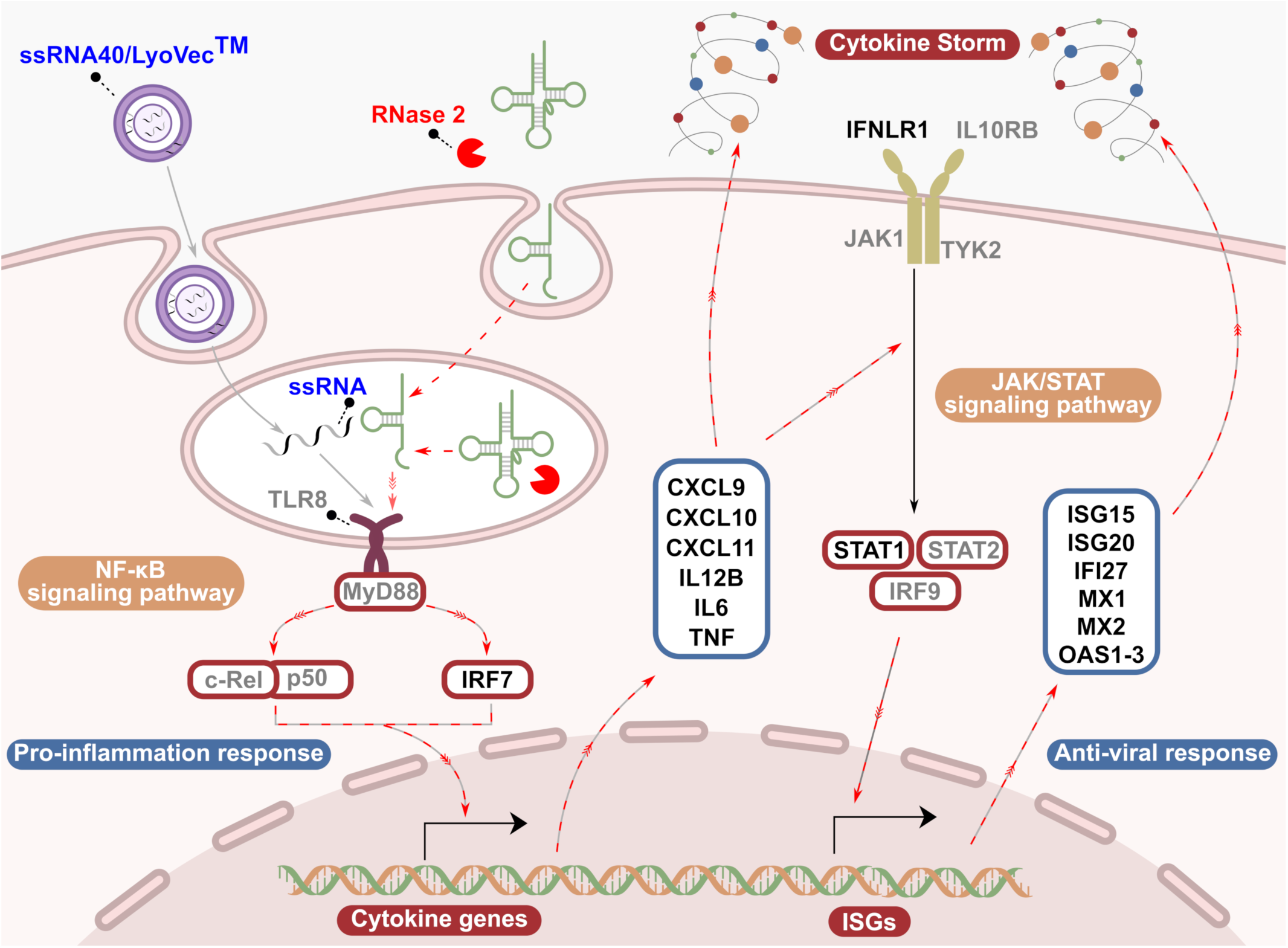
Schematic illustration of the proposed molecular mechanism of RNase 2 in macrophages when exposed to viral RNA. The most significant changed genes associated to RNase 2 and ssRNA40 exposure by IR analysis are in black (cut-off: log_2_FC>5, padj<0.01) and grey (cut-off: log_2_FC>0, padj<0.01). The pathways where RNase 2 may be involved are indicated by red dashed lines overlapping with standard pathways of immune response to viral RNA exposure.

Our previous work corroborated the selective cleavage on ncRNAs by RNase 2 in THP1-derived macrophages^15^ using synthetic tRNAs and anticodon loop hairpins.^36^ The current work adds a 3^rd^ layer of information by an integrative transcriptome analysis of mRNA-small RNA population in basal and cell activation by viral RNA conditions. We have compared 3 types of small RNA population libraries according to their nucleic acid ends including amplification of RNA products generated by an endonuclease cleavage. Notwithstanding, we are aware that NGS methodology can be biased by the presence of posttranscriptional modifications. Thus, future work should benefit from an epitranscriptomic analysis together with additional information on RNA structuration and potential ribonucleoprotein complex formation *in vivo* that will determine the RNase accessibility to its target.^90^ Besides, to obtain a full account of RNase 2 activity, current studies on macrophages need to be expanded to other more complex cellular models, in a physiological relevant environment.

## ACKNOWLEDGMENTS

We thank the Spanish Agencia Estatal de Investigación for funding (PID2022-137872NB-I00).

## MATERIALS and METHODS

### Cell culture and treatment

The Human THP-1 RNase 2-knockout (KO) cells were obtained as described before.^15^ Human THP-1 wild-type (WT) and RNase 2-knockout (KO) cells were maintained or passaged in 25 or 75 cm^2^ tissue culture flasks using RPMI-1640 medium supplied with 10% heat-inactivated FBS at 37°C in humidified 5% CO_2_ conditions. When the desired number of cells has reached, THP-1 cells were then induced to macrophage by 50 nM PMA for 48h as previously described.^15^ Total 10^6^ cells/well were seeded in 6-well cell culture plates. Four groups have been set up, which correspond to THP-1 WT (R2WT), THP-1 KO (R2KO), THP-1 WT plus 4 μg/ml ssRNA40 (WT40) and THP-1 KO plus 4 μg/ml ssRNA40 (KO40) conditions. The ssRNA40 was complemented with LyoVec™ (InvivoGen) transfection reagent. After 14h incubation with/without ssRNA40, cells were first washed with cold PBS and then RNA was extracted using the methods described below.

### RNA extraction and library preparation

The membrane of attached macrophage cells in the 6-well plate was broken using 600 μl lysis buffer from

*mir*Vana^TM^ miRNA isolation kit (Invitrogen™, AM1561). Total 4 libraries were prepared (Figure S7).

► **Library 1:** mRNA was extracted as instructed in the *mir*Vana^TM^ miRNA isolation kit for total RNA isolation procedure.

Three small RNA libraries were prepared according to the different end of small RNA.

► **Library 2**: small RNA (<200nt) was obtained directly by using *mir*Vana^TM^ miRNA isolation kit to enrich small RNA content.

► **Library 3:** small RNA (<200nt) was firstly obtained by using *mir*Vana^TM^ miRNA isolation kit as Library 2. 20 to 100 nt RNAs were then extracted from 8% TBE-PAGE gel. By following the previous method,^15,35^ the small RNA was purified only ending with cP in 25-100 nt. Briefly, the purified RNAs were firstly treated by calf intestinal alkaline phosphatase (CIP, New England Biolabs, M0525S). After phenol–chloroform purification, the RNAs were oxidized by incubation in 10 mM NaIO_4_ (Sigma-Aldrich, 311448) at 0 °C for 40 min in the dark, followed by ethanol precipitation. The RNAs were finally treated with T4 Polynucleotide Kinase (T4 PNK, Thermo Scientific, EK0031), followed by another phenol–chloroform purification. Phosphatase treatment with CIP removes phosphate group at either 5’ or 3’ end. Following, periodate cleaves the cis-diol group of 3’OH end to generate 2’,3’-dialdehydes that could not ligate with adaptors for the amplification step, leaving only the RNA molecules with a 2’3’cyclic phosphate linkage.^35^

► **Library 4**: small RNA (<200nt) was firstly obtained by using *mir*Vana^TM^ miRNA isolation kit as Library 2 and then processing as Library 3 by skipping the NaIO_4_ treatment.

RNA purity was determined by spectrophotometry and RNA integrity was analyzed using Agilent 2100 Bioanalyzer and calculated as an RNA integrity number. Following RNA extraction, different RNA samples were submitted to CRG genomics sequencing service (Centre for Genomic Regulation, Barcelona, Spain) for cDNA library preparation and NGS sequencing. Sequencing libraries were prepared according to protocols provided by Illumina. 50bp-long paired end sequencing for Library 1 with a depth of > 50 million reads per sample and 150bp-long paired end sequencing for Library 2-4 with a depth of > 30 million reads per sample were carried out in an Illumina NovaSeq6000 platform.

### Data processing

► **Library 1**: For mRNA processing, fastp (v0.23.4)^91^ was used to trim raw sequencing data and discard low-quality reads (Q<20). The trimmed pair-end read transcript sequences were then aligned to reference genome (GRCh38.primary_assembly.genome, Gencode.v47.transcripts) through salmon (v1.10.1)^92^ and samtools (v1.9)^93^ was used to sort the sam to bam files. The raw count was exported by tximport R package (v1.30.0)^94^ and differential expression analysis was executed by DESeq2 (v1.42.0) R package among groups.^33^ Biomart R package tool (v2.58.0) was used to classify the transcripts according to their biotype.^95^

► **Library 2-4**: For ncRNA analysis, overall biotypes in each ncRNA library have been firstly annotated. Briefly, the pair-end raw reads with adapters were firstly trimmed and filtered (Q>20) by Cutadapt (v4.8)^96^ and both end reads, merged by Seqprep (https://github.com/jstjohn/SeqPrep, v1.2), were output with two different cut-off of length (reads length ≤ 50nt and reads length > 50nt). Subsequently, the merged reads were mapped to human genome (GRCh38.primary_assembly.genome) integrated with ssRNA40 fragment through Bowtie 1 (v1.3.1)^97^ for the short reads (length ≤ 50nt) and Bowtie2 (v2.5.1)^98^ for the long reads (length > 50nt) separately. The aligned reads were then counted and annotated by feacture.count^99^ according to the reference GTF files available on the gencode website (gencode.v47.primary_assembly.annotation.gtf and gencode.v47.tRNAs.gtf) and piRNA database (piRNAdb: piRNAdb.hsa.v1_7_6.hg38.fa).

For **miRNA analysis** in **Library 2**, the merged fastq files above were applied into miRDeep2 (v0.1.3)^100^ to identify and count the miRNA (miRbase, release 22.1) by following the method described previously.^101^ In short, *mapper.pl* was used to process all the fastq reads and generate the input files for *miRDeep2.pl*. The *hairpin.fa* and *mature.fa* files from miRBase database (https://www.mirbase.org/download/) were used to annotate the identified miRNAs. The final integrated count file from *quantifier.pl* including miRNA (total read count > 20) was used for differential expression analysis with DESeq2 (v1.42.0).

For **tRNA analysis** of 2’3’cP RNA-seq in **Library 3**, the trimmed and merged fastq files above were aligned to the hg38 by following the standard workflows of tRAX pipeline (https://trna.ucsc.edu/tRAX/).^102^ Briefly, the reference database was built by *maketrnadb.py* with hg38 from UCSC human genome and GtRNAdb database. Following, *processamples.py* was executed to map and annotate the tRNA and tDRs. Unique raw count of tRNA&tDRs (total read count > 20) were then used for downstream analyses by DEseq2 (v1.42.0). The produced bam files were visualized by Integrative Genomics Viewer (IGV) to check the enriched tDRs.

### Data analysis and visualization

Interaction analysis (IR analysis) was conducted through DESeq2 (v1.42.0) by ∼genotype + condition + genotype:condition model.^33^ GO enrichment and KEGG pathway enrichment analysis of the differential expression of genes across the samples was carried out using the clusterProfiler R package (v4.10.0).^103^ WikiPathway analysis was based on the results obtained from g: Profiler by using a 0.05 threshold (https://biit.cs.ut.ee/gprofiler/gost). R packages including ggplot2 (v3.5.0),^104^ EnhancedVolcano (v1.20.0, https://github.com/kevinblighe/EnhancedVolcano), simplifyEnrichment (v1.12.0),^105^ ComplexHeatmap (v2.18.0)^106^ and Pathview (v3)^107^ were installed in Conda (v23.7.3) environments to visualize the results.

