## Supplementary Material for "Human RNase 2 is essential for macrophage response to viral RNA"

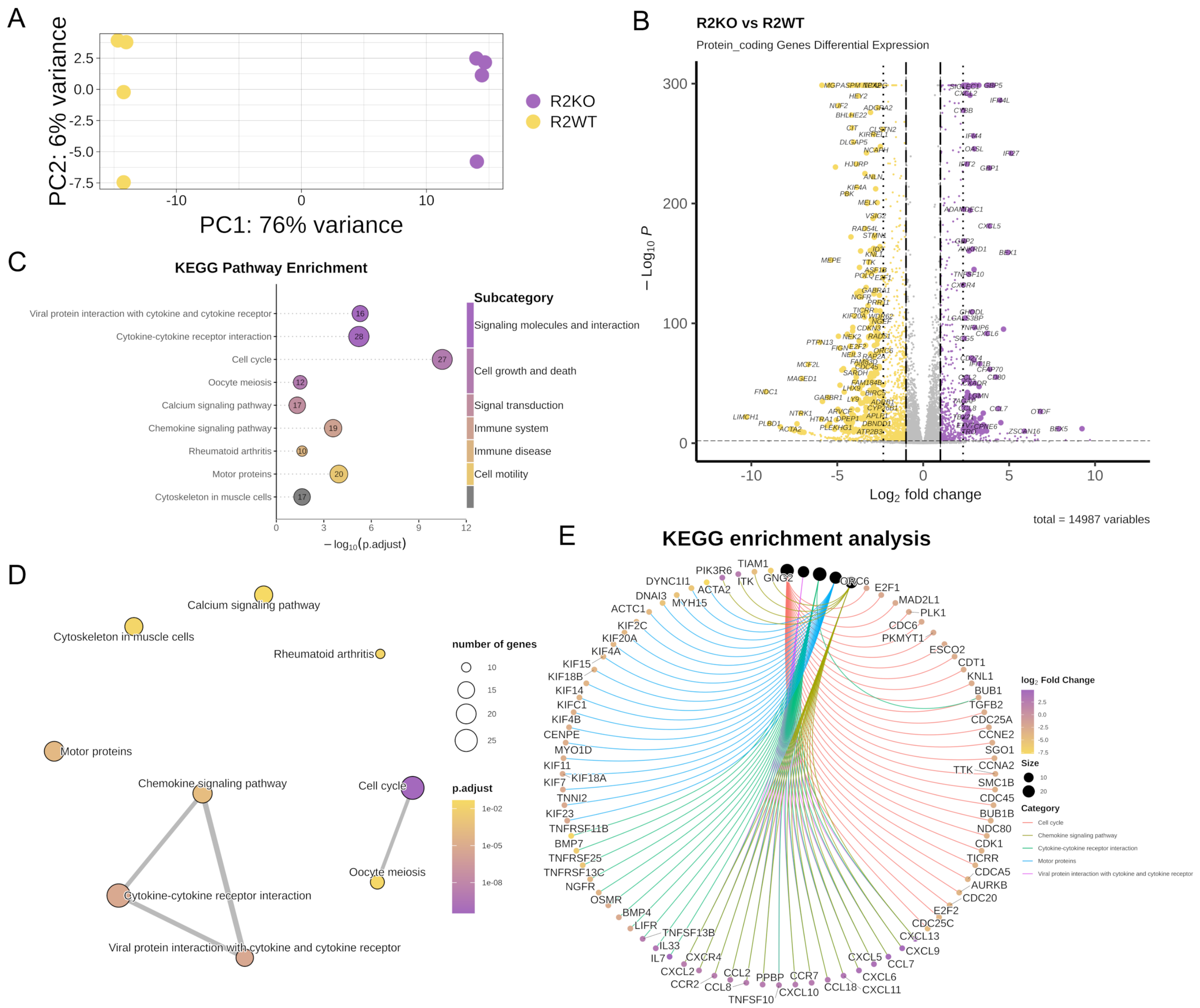


**Supplemental Figure S1. Overview of the results of 1st NGS for RNase 2-KO (R2KO) and WT (R2WT) THP-1 induced macrophages.**

**(A)** Principal Component Analysis (PCA) plot for R2WT and R2KO macrophage cells of poly(A) transcripts' sequencing.

**(B)** Volcano plot for protein_coding transcripts in R2KO vs R2WT cells. The significantly down-regulated and up-regulated protein coding genes identified from NGS are shown (|$Log_2$FoldChange|>1 and padj<0.01) and the top significant transcripts (|$Log_2$FoldChange|>2.32 and padj<$10^{-10}$) are labelled with their name.

**(C)** KEGG pathway enrichment dotplot of R2KO vs R2WT. The significantly down-regulated and up-regulated protein coding genes for R2KO vs R2WT (|$Log_2$FoldChange|>2.32 and padj<0.01) are selected to plot the figure. The identified significantly KEGG pathways (qvalue<0.05) with the number of genes inside the corresponding dots of each item are shown. The subcategories of each item in KEGG database (https://www.genome.jp/kegg/pathway.html) are summarized on the right side.

**(D)** KEGG pathway enrichment emapplot of R2KO vs R2WT. The significantly down-regulated and up-regulated protein coding genes for R2KO vs R2WT (|Log2FoldChange|>2.32 and padj<0.01) are selected.

**(E)** KEGG pathway enrichment cnetplot of R2KO vs R2WT. The top 5 pathways for the category of environmental information processing in KEGG database were plotted. The $log_2$Fold Change (R2KO vs R2WT) of protein coding genes was shown with a color gradient for up to down.

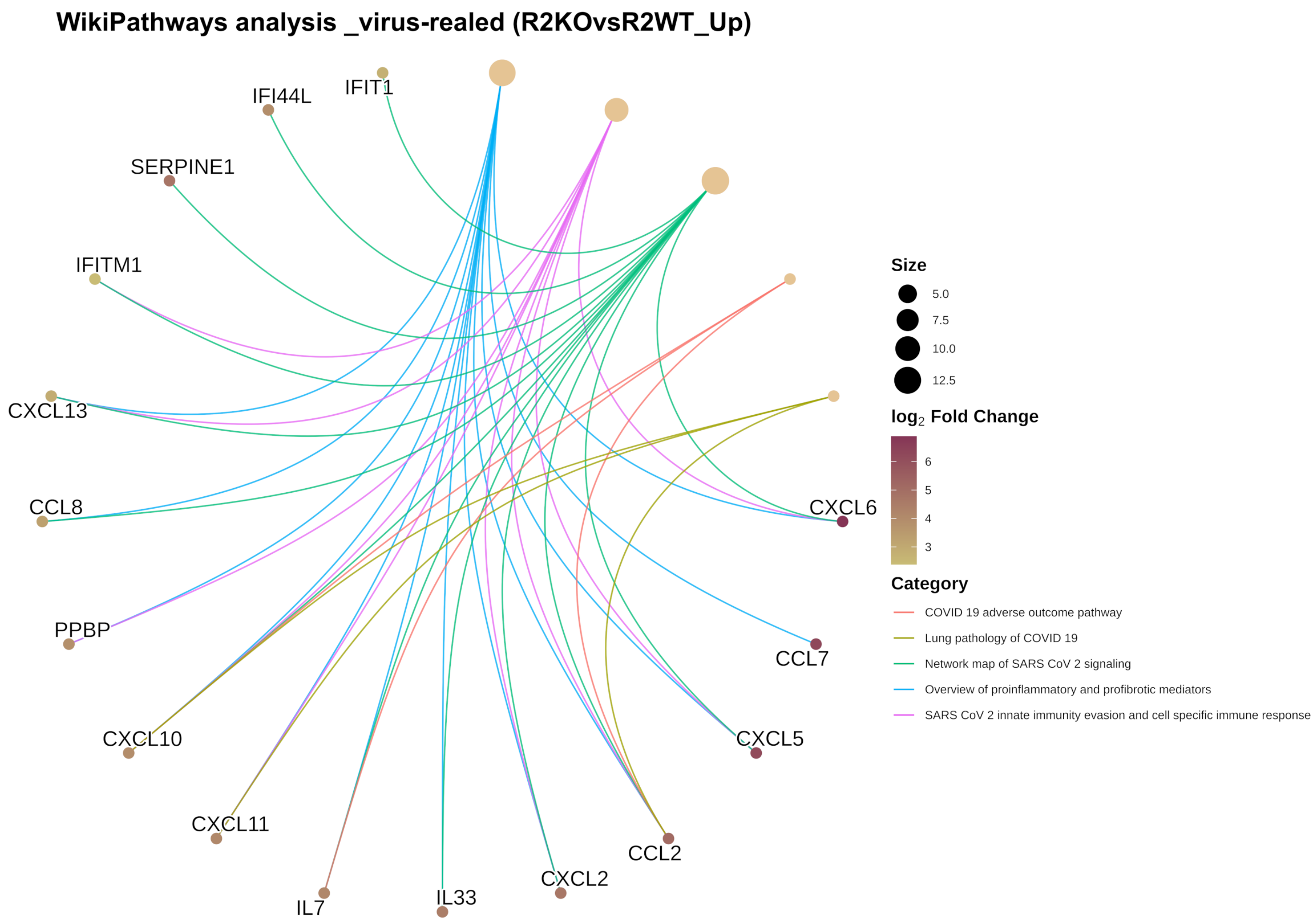


**Supplemental Figure S2. Wikipathway enrichment cnetplot of R2KO vs R2WT.** The significantly up-regulated protein coding genes for R2KO vs R2WT (|$Log_2$FoldChange|>2.32 and padj<0.01) from 1st and 2nd NGS were selected to plot the figure. The significant pathways (qvalue<0.05) are obtained from g:Profiler database (https://biit.cs.ut.ee/gprofiler/gost, access date: 05/09/2025) by uploading the official gene names.

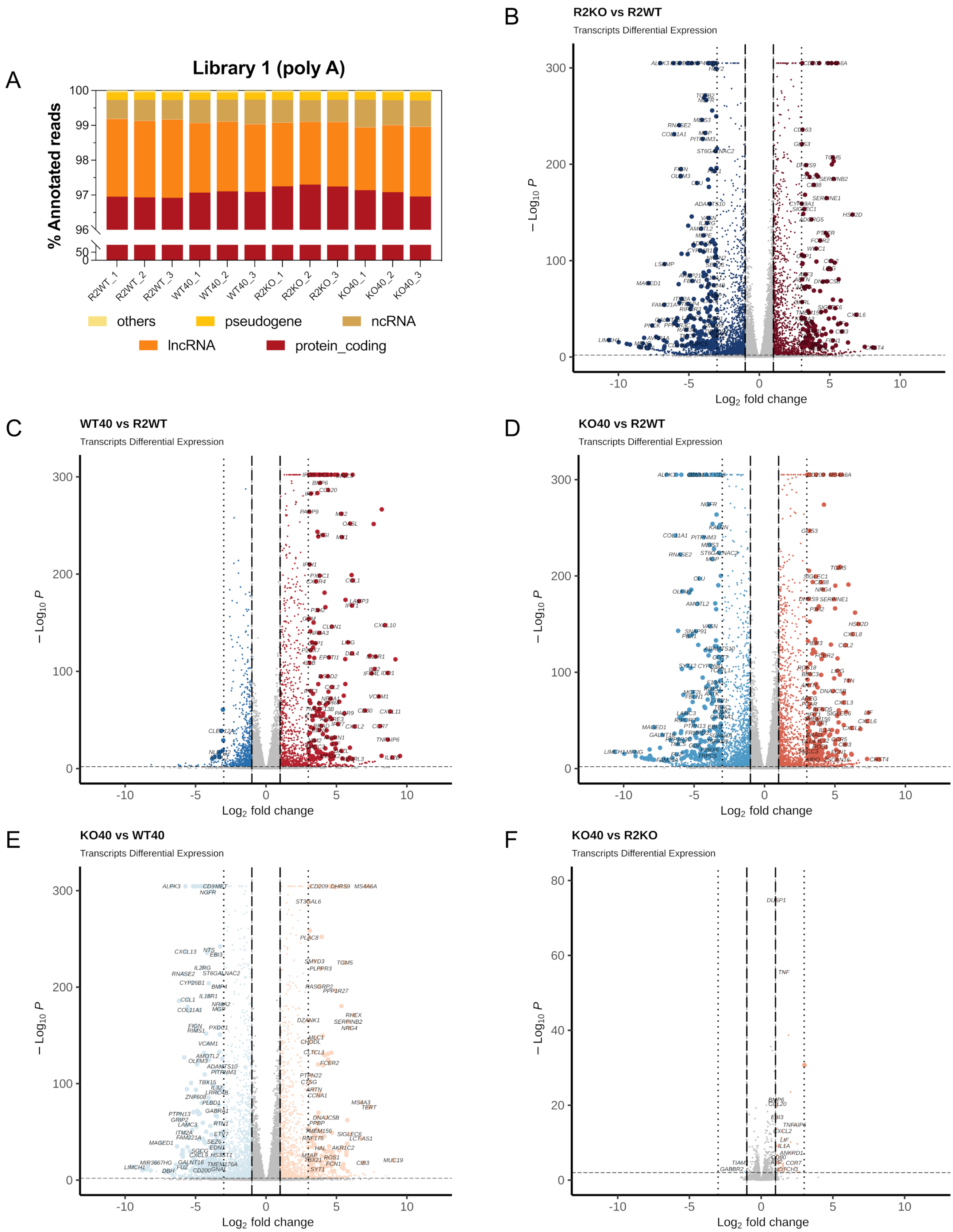


**Supplemental Figure S3. Overview of the results of 2nd NGS for RNase 2-KO (R2KO) and WT (R2WT) THP-1 induced macrophage in absence/presence of ssRNA40.**

**(A)** Poly(A) transcript types were identified by alignment with the human genome.

**(B-F)** Volcano plot for transcripts with different two-by-two comparison. The significantly down-regulated and up-regulated transcripts identified from NGS are shown ($|Log_2FoldChange|>1$ and $padj<0.01$) and the top significant transcripts ($|Log_2FoldChange|>3$ and $padj<10^{-10}$) are labelled with their name.

R2WT is the control where the standard THP-1 induced macrophages without any treatment were used. WT40 is the R2WT cells treated with 4 µg/ml ssRNA40 for 14 hours. R2KO/KO40 corresponds to the KO macrophage cells in absence/presence of ssRNA40.

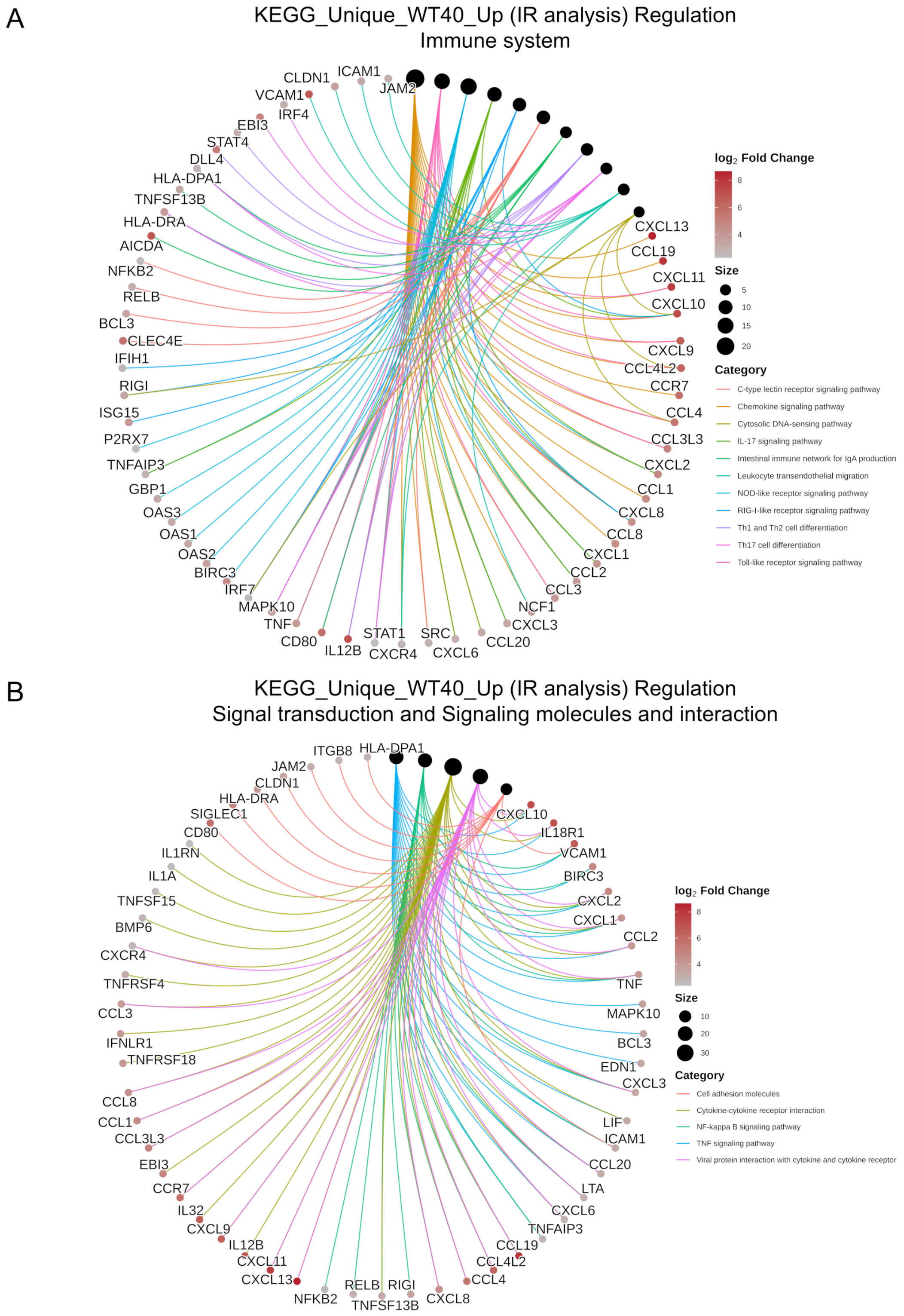


**Supplemental Figure S4. Genes involved in ssRNA40 activation related to RNase 2.**
**(A)** Cnetplot of KEGG pathways for immune system subcategory.
**(B)** Cnetplot of KEGG pathways for signal transduction and signaling molecules and interaction subcategory.

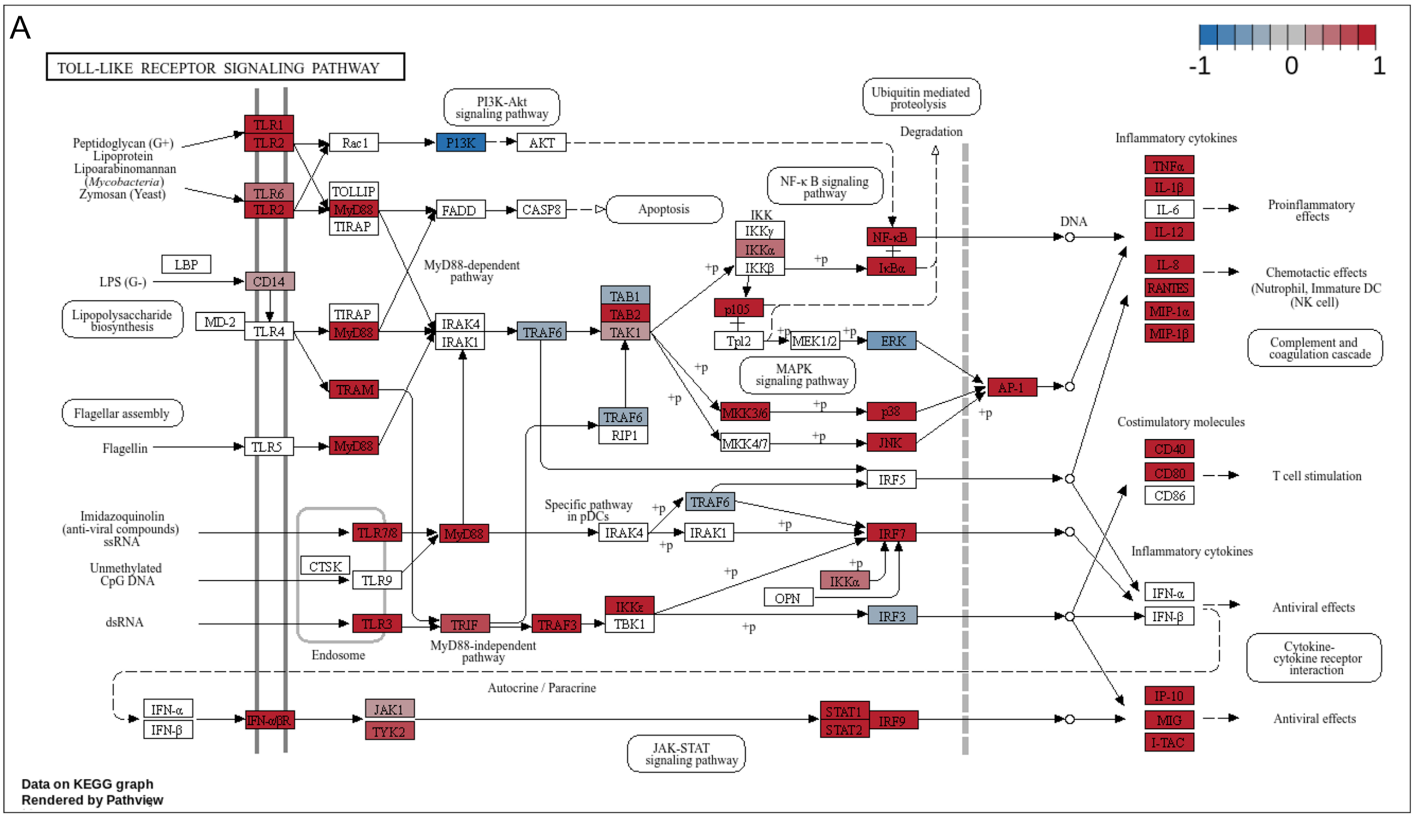
A
TOLL-LIKE RECEPTOR SIGNALING PATHWAY
-1
0
1
Peptidoglycan (G+)
Lipoprotein
Lipoarabinomannan
(Mycobacteria)
Zymosan (Yeast)
TLR1
TLR2
TLR6
TLR2
Rac1
PI3K
AKT
PI3K-Akt signaling pathway
TOLLIP
MyD88
TIRAP
FADD
CASP8
Apoptosis
LBP
LPS (G-)
CD14
Lipopolysaccharide biosynthesis
MD-2
TLR4
TIRAP
MyD88
TRAM
MyD88-dependent pathway
IRAK4
IRAK1
TRAF6
TAB1
TAB2
TAK1
TRAF6
RIP1
Flagellar assembly
Flagellin
TLR5
MyD88
IKK
IKKγ
IKKα
IKKβ
NF-κB signaling pathway
NF-κB
IκBα
p105
Tpl2
MEK1/2
ERK
MAPK signaling pathway
MKK3/6
MKK4/7
p38
JNK
AP-1
Ubiquitin mediated proteolysis
Degradation
DNA
+p
Inflammatory cytokines
TNFα
IL-1β
IL-6
IL-12
Proinflammatory effects
IL-8
RANTES
MIP-1α
MIP-1β
Chemotactic effects
(Nutrophil, Immature DC
(NK cell)
Complement and coagulation cascade
Costimulatory molecules
CD40
CD80
CD86
T cell stimulation
IRF5
Imidazoquinolin
(anti-viral compounds)
ssRNA
Unmethylated
CpG DNA
dsRNA
CTSK
TLR7/8
TLR9
TLR3
Endosome
MyD88
Specific pathway in pDCs
IRAK4
IRAK1
TRAF6
IRF7
IKKα
OPN
TRIF
TRAF3
IKKε
TBK1
IRF3
MyD88-independent pathway
Inflammatory cytokines
IFN-α
IFN-β
Antiviral effects
Cytokine-cytokine receptor interaction
Autocrine / Paracrine
IFN-α
IFN-β
IFN-α/βR
JAK1
TYK2
STAT1
STAT2
IRF9
JAK-STAT signaling pathway
IP-10
MIG
I-TAC
Antiviral effects
Data on KEGG graph
Rendered by Pathview

B

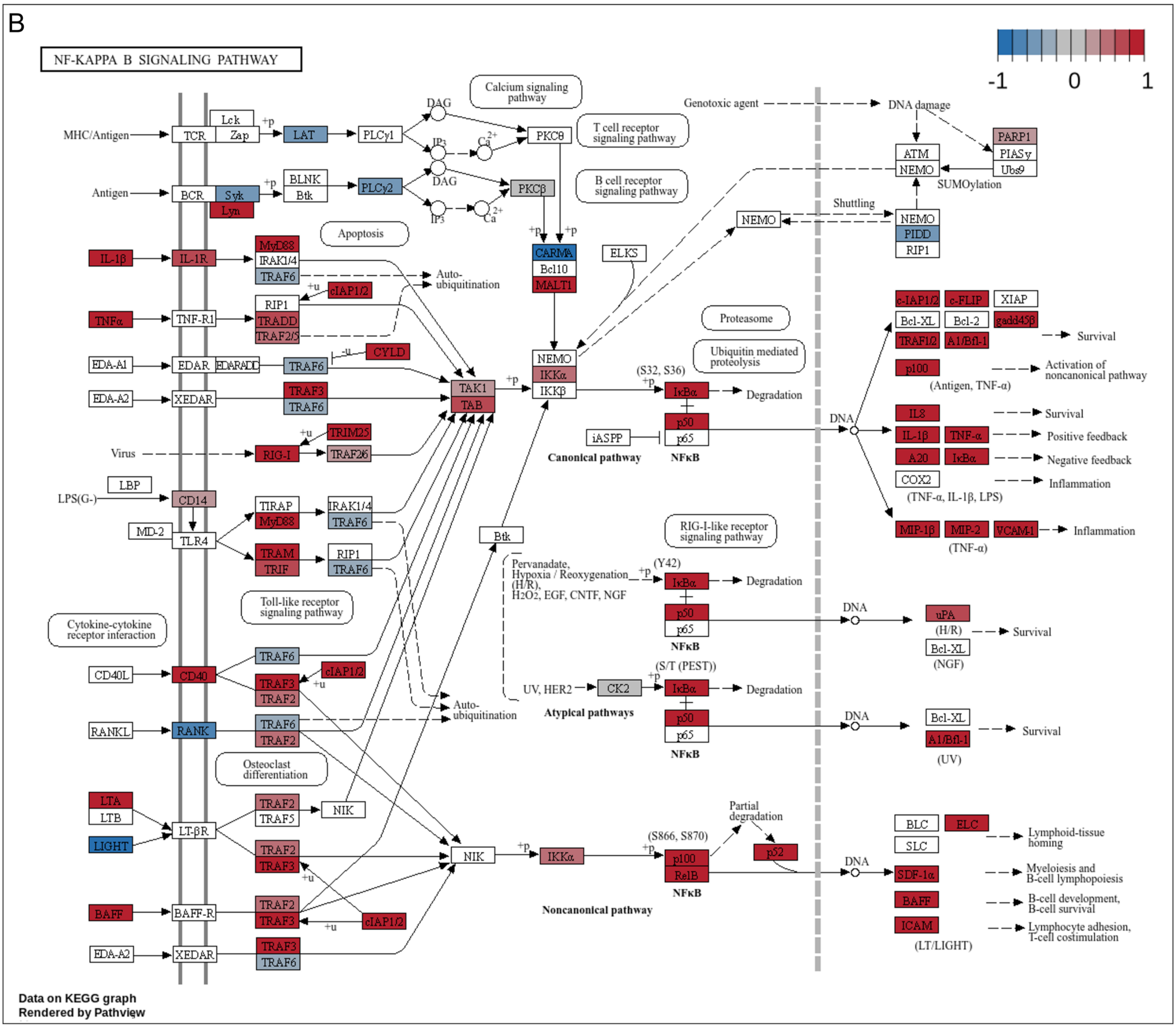
NF-KAPPA B SIGNALING PATHWAY
-1
0
1
MHC/Antigen
TCR
Lck
Zap
LAT
PLCγ1
DAG
IP3
PKCθ
Calcium signaling pathway
T cell receptor signaling pathway
Antigen
BCR
Syk
Lyn
BLNK
Btk
PLCγ2
PKCβ
B cell receptor signaling pathway
Apoptosis
IL-1β
IL-1R
MyD88
IRAK1/4
TRAF6
cIAP1/2
Auto-ubiquitination
TNFα
TNF-R1
RIP1
TRADD
TRAF2/5
CYLD
EDA-A1
EDAR
EDARADD
EDA-A2
XEDAR
TRAF3
TAK1
TAB
CARMA
Bcl10
MALT1
ELKS
NEMO
IKKα
IKKβ
Virus
RIG-I
TRIM25
TRAF26
LBP
LPS(G-)
CD14
MD-2
TLR4
TIRAP
TRAM
TRIF
Btk
Toll-like receptor signaling pathway
Cytokine-cytokine receptor interaction
CD40L
CD40
RANKL
RANK
Osteoclast differentiation
LTA
LTB
LIGHT
LT-βR
TRAF5
NIK
BAFF
BAFF-R
Genotoxic agent
DNA damage
ATM
PARP1
PIASy
Ubc9
SUMOylation
Shuttling
PIDD
Proteasome
Ubiquitin mediated proteolysis
(S32, S36)
IκBα
Degradation
p50
p65
iASPP
Canonical pathway
NFκB
DNA
c-IAP1/2
c-FLIP
XIAP
Bcl-XL
Bcl-2
gadd45β
TRAF1/2
A1/Bfl-1
Survival
p100
Activation of noncanonical pathway
(Antigen, TNF-α)
IL8
IL-1β
TNF-α
A20
COX2
Positive feedback
Negative feedback
Inflammation
(TNF-α, IL-1β, LPS)
MIP-1β
MIP-2
VCAM-1
(TNF-α)
RIG-I-like receptor signaling pathway
Pervanadate, Hypoxia / Reoxygenation (H/R), H2O2, EGF, CNTF, NGF
(Y42)
uPA
(H/R)
(NGF)
UV, HER2
CK2
(S/T (PEST))
Atypical pathways
(UV)
Partial degradation
(S866, S870)
RelB
p52
Noncanonical pathway
BLC
ELC
SLC
SDF-1α
ICAM
(LT/LIGHT)
Lymphoid-tissue homing
Myeloiesis and B-cell lymphopoiesis
B-cell development, B-cell survival
Lymphocyte adhesion, T-cell costimulation
Data on KEGG graph
Rendered by Pathview

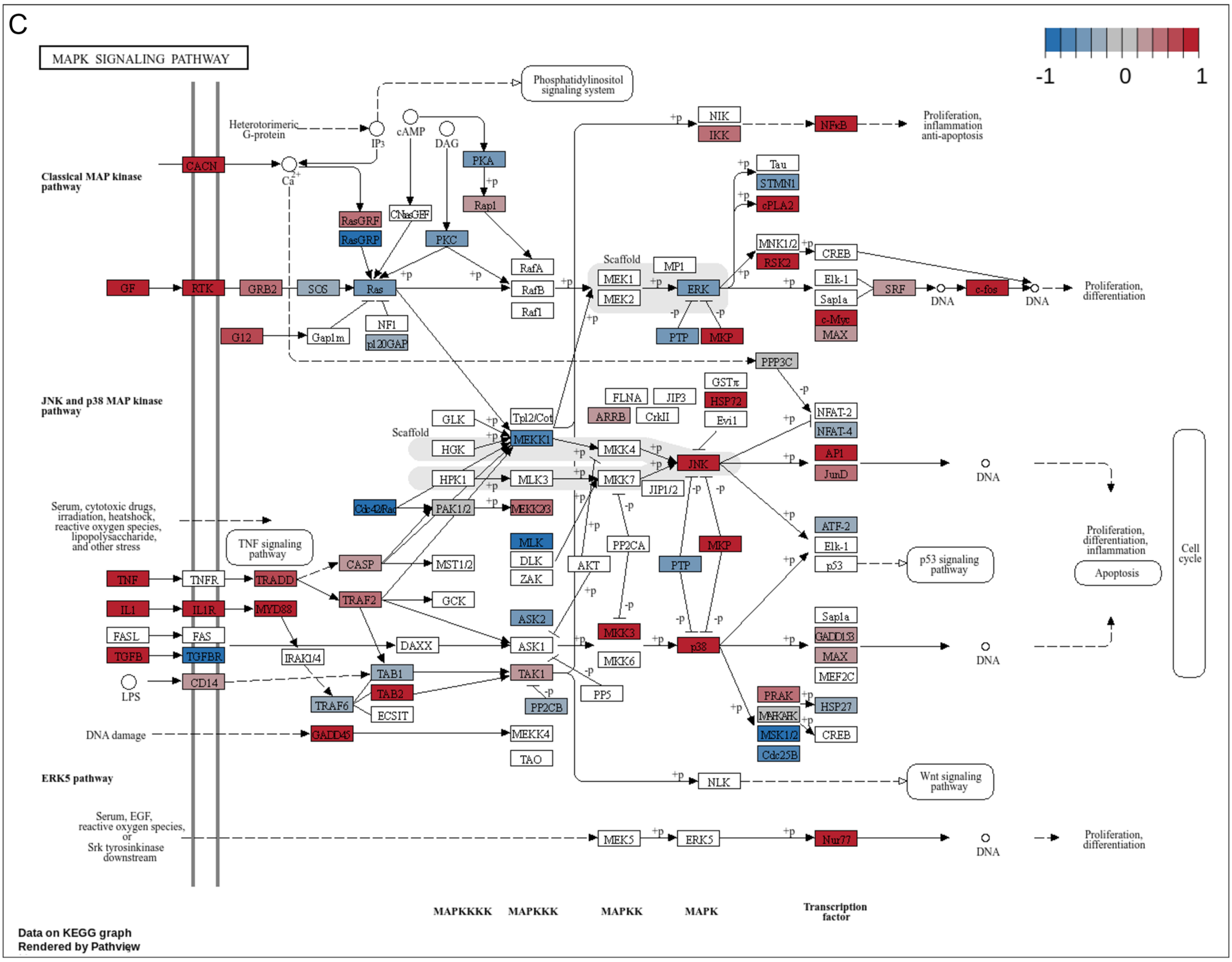
C
MAPK SIGNALING PATHWAY
-1
0
1
Phosphatidylinositol signaling system
Heterotrimeric G-protein
IP3
cAMP
DAG
Classical MAP kinase pathway
CACN
PKA
Rap1
RasGRF
RasGRP
CNrasGEF
PKC
GF
RTK
GRB2
SOS
Ras
RafA
RafB
Raf1
NF1
p120GAP
G12
Gap1m
Scaffold
MEK1
MEK2
MP1
ERK
PTP
MKP
NIK
IKK
NFκB
Proliferation, inflammation anti-apoptosis
Tau
STMN1
cPLA2
MNK1/2
RSK2
CREB
Elk-1
Sap1a
SRF
c-fos
c-Myc
MAX
DNA
Proliferation, differentiation
PPP3C
JNK and p38 MAP kinase pathway
GSTπ
HSP72
FLNA
JIP3
ARRB
CrkII
Evi1
NFAT-2
NFAT-4
AP1
JunD
GLK
Tpl2/Cot
MEKK1
HGK
HPK1
MLK3
MKK4
MKK7
JIP1/2
JNK
Cdc42/Rac
PAK1/2
MEKK2/3
Serum, cytotoxic drugs, irradiation, heatshock, reactive oxygen species, lipopolysaccharide, and other stress
TNF signaling pathway
CASP
MST1/2
MLK
DLK
ZAK
AKT
PP2CA
PTP
MKP
ATF-2
Elk-1
p53
p53 signaling pathway
Proliferation, differentiation, inflammation
Apoptosis
Cell cycle
TNF
TNFR
TRADD
IL1
IL1R
MYD88
TRAF2
GCK
ASK2
FASL
FAS
TGFB
TGFBR
IRAK1/4
DAXX
ASK1
MKK3
MKK6
p38
Sap1a
GADD153
MAX
MEF2C
LPS
CD14
TAB1
TAB2
TRAF6
ECSIT
TAK1
PP2CB
PP5
PRAK
MAPKAPK
MSK1/2
Cdc25B
HSP27
CREB
DNA damage
GADD45
MEKK4
TAO
ERK5 pathway
NLK
Wnt signaling pathway
Serum, EGF, reactive oxygen species, or Srk tyrosinkinase downstream
MEK5
ERK5
Nur77
MAPKKKK
MAPKKK
MAPKK
MAPK
Transcription factor
Data on KEGG graph
Rendered by Pathview

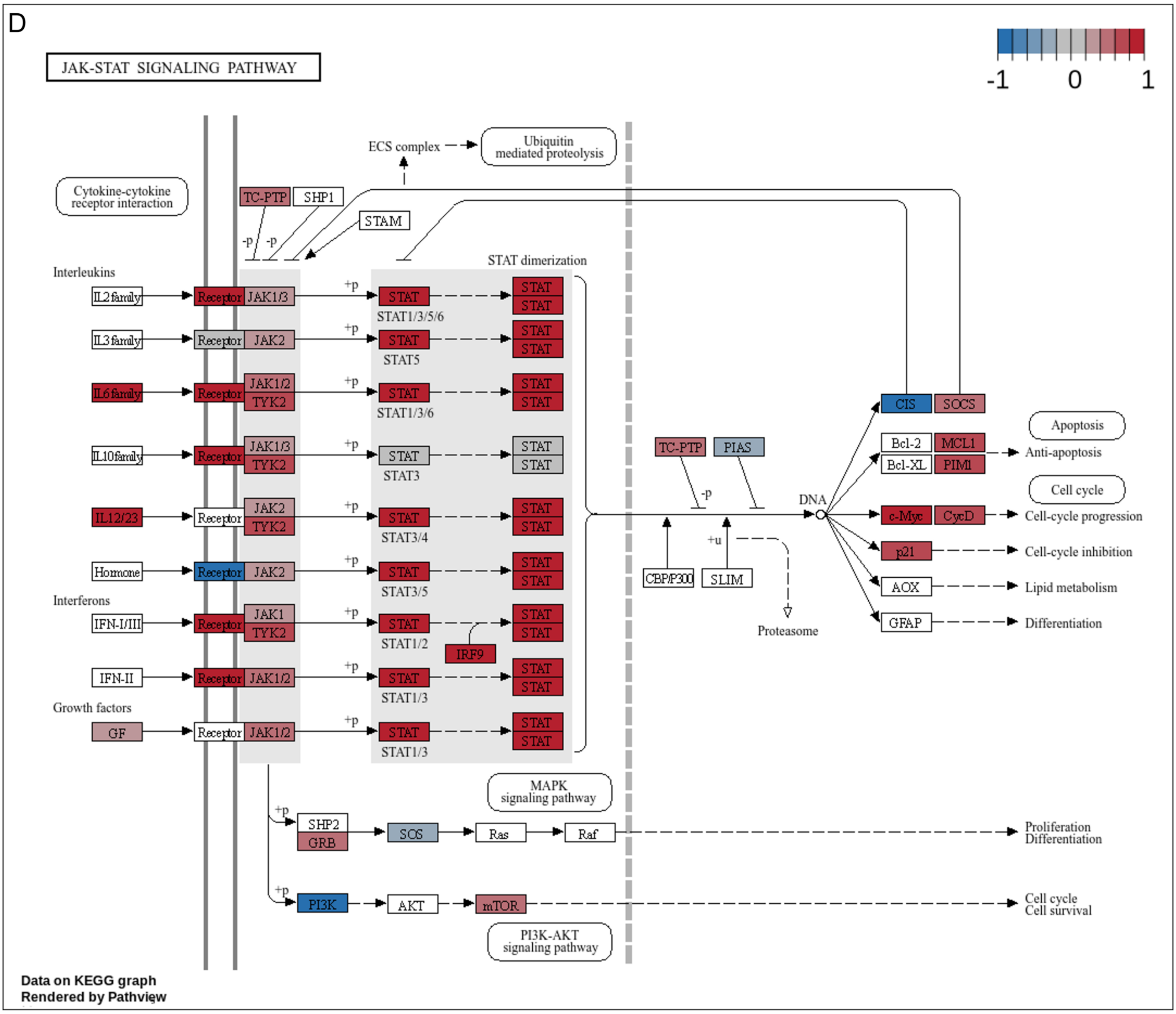


**Supplemental Figure S5**. **Pathview of selected KEGG enrichment in WT40 subgroup.**

(A) Pathview of TLR-like receptor signaling pathway (hsa04620)

(B) Pathview of NF-κB signaling pathway (hsa04064)

(C) Pathview of MAPK signaling pathway (hsa04010)

(D) Pathview of JAK-STAT signaling pathway (hsa04630)

The filter genes that related to RNase 2 upon ssRNA40 exposure after interaction analysis were colored in the figures where red indicates up-regulation and blue down-regulation.

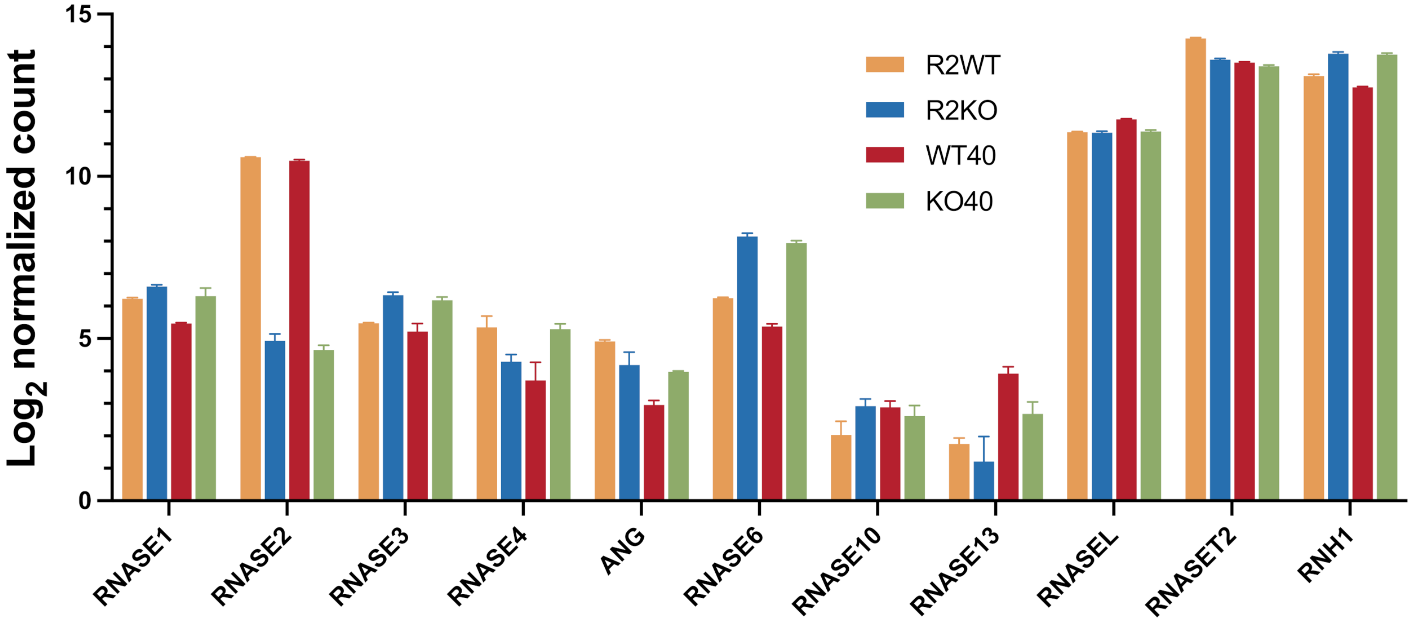


**Supplemental Figure S6. Log$_2$ normalized count from NGS analysis corresponding to expression of RNase A family members, other selected RNases, and RNase inhibitor (RNH1) for each group condition of THP-1-derived macrophages.** R2WT is the control where the standard THP-1 induced macrophages without any treatment were used. WT40 is the R2WT cells treated with 4 µg/ml ssRNA40 for 14 hours. R2KO/KO40 corresponds to the KO macrophage cells in absence/presence of ssRNA40.

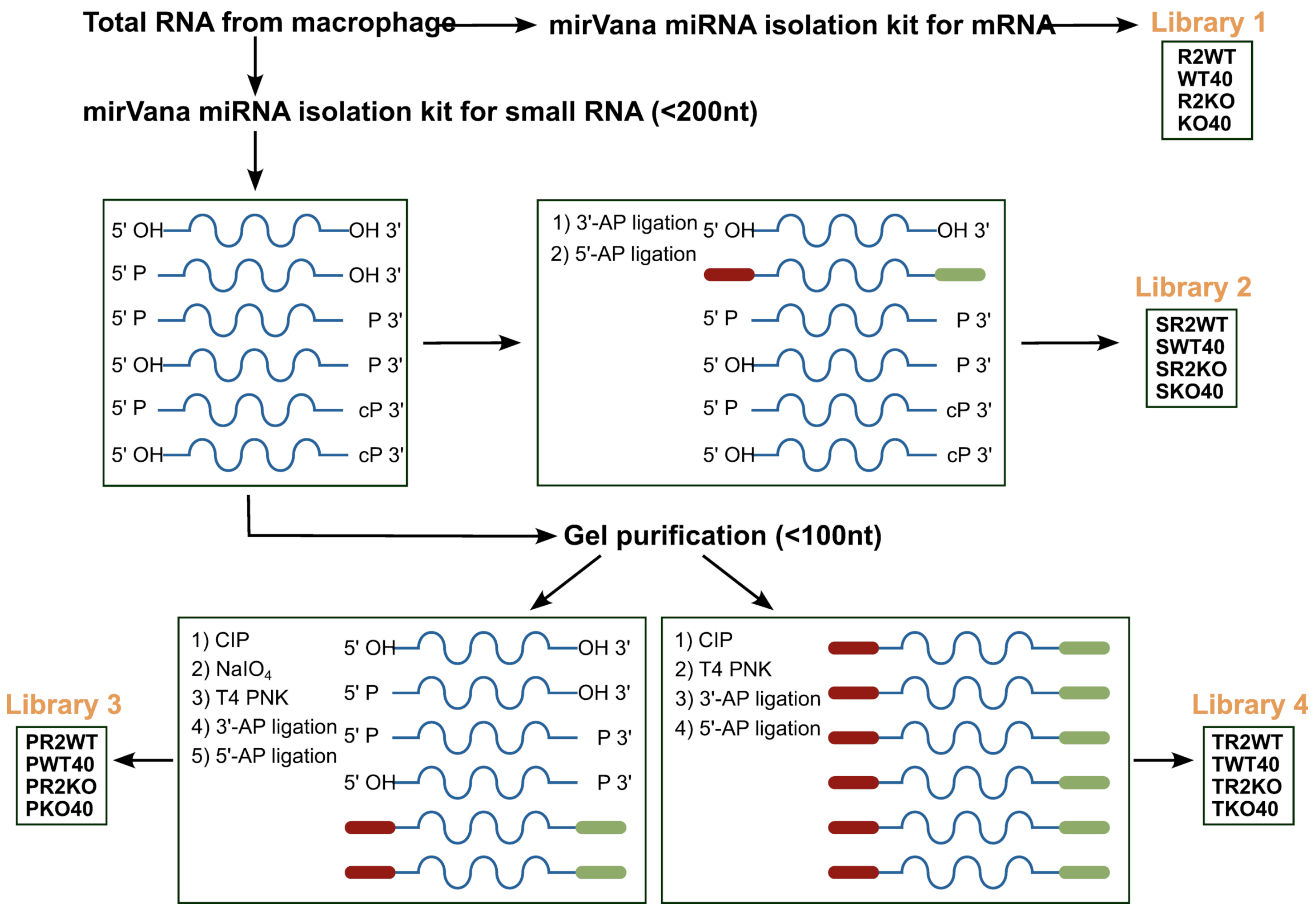


**Supplemental Figure S7. Scheme of RNA libraries preparation.** 4 libraries were obtained with different treatments and correspond to next-generation sequencing (NGS) of: mRNA (Library 1), standard small RNAs starting with 5'phosphate (5'P) and ending with 3' hydroxyl (3'OH) (Library 2), selective amplification of 2'3' cyclic phosphate (3'cP) end products (Library 3), which are generated by an endonuclease cleavage, and total small RNA with any ends (Library 4). R2WT is the control corresponding to THP-1-derived macrophages without any treatment. WT40 is the R2WT cells treated with 4 µg/ml ssRNA40 for 14 hours. R2KO/KO40 corresponds to the KO macrophage cells in absence/presence of ssRNA40.

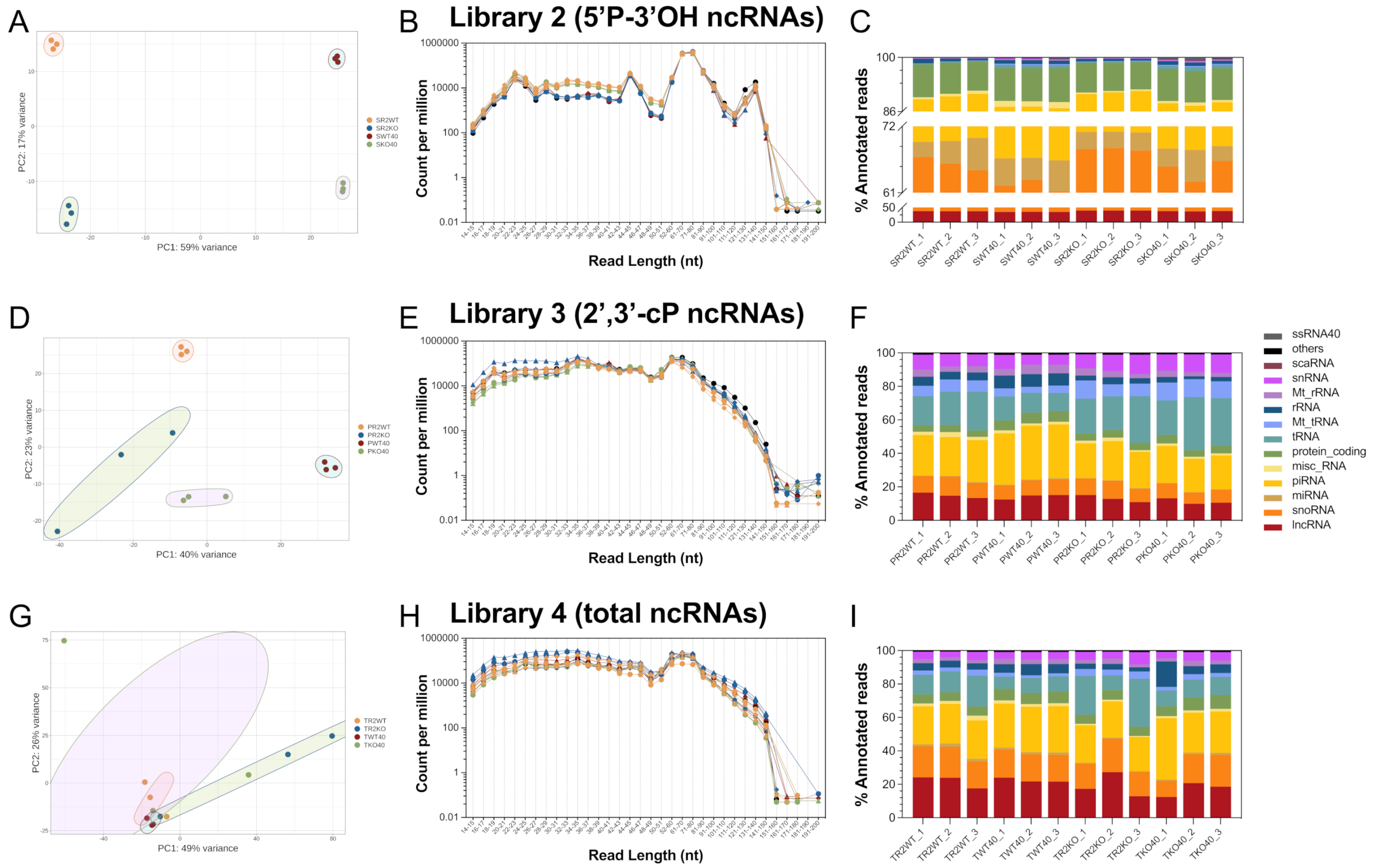


**Supplemental Figure S8. Overview of prepared ncRNA libraries**.

**(A-C)** Library 2 for standard ncRNAs with 5'P and 3'OH.

**(D-F)** Library 3 for ncRNAs harboring with 3'cP end.

**(G-I)** Library 4 for ncRNAs with any end.

Libraries preparation is described in Figure S7. On the left, for each library genome we include Principal Component Analysis (PCA) plot, read counts distributed by length and RNA types annotated to the human for 4 groups.

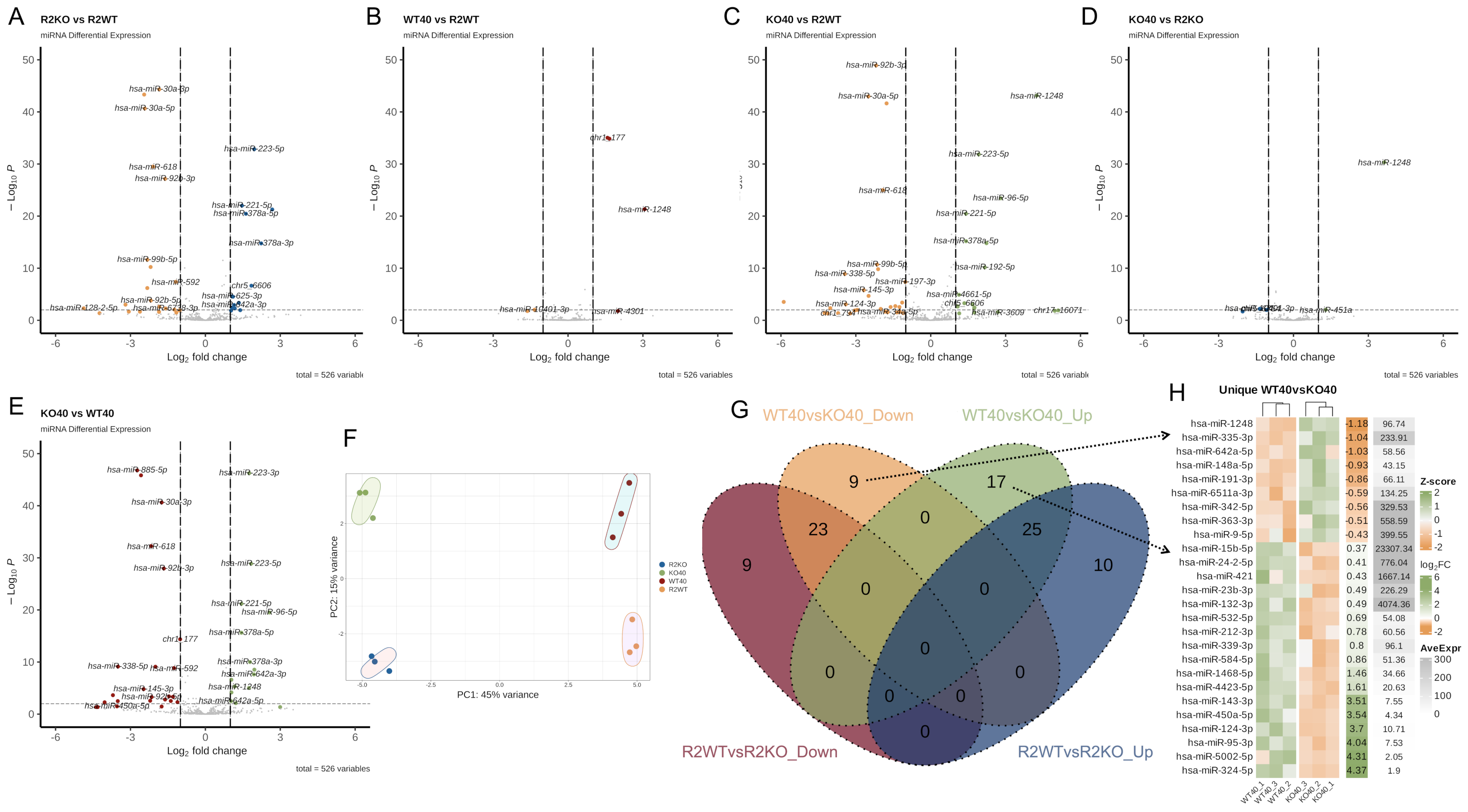


**Supplemental Figure S9. miRNA comparison**.

**(A-E)** Volcano plot for transcripts with different two-by-two comparison**.** The significantly down/up-represented miRNAs identified from miRDeep2 are labelled with the miRBase names (Cut-off: $|Log_2FC|>1$ and padj<0.01).

**(F)** PCA plot of miRNAs among 4 group.

**(G)** Venn plot of up/down-represented miRNAs for WT40 vs KO40 and R2WT vs R2KO.

**(H)** Heatmap for unique miRNAs in WT40 vs KO40 group

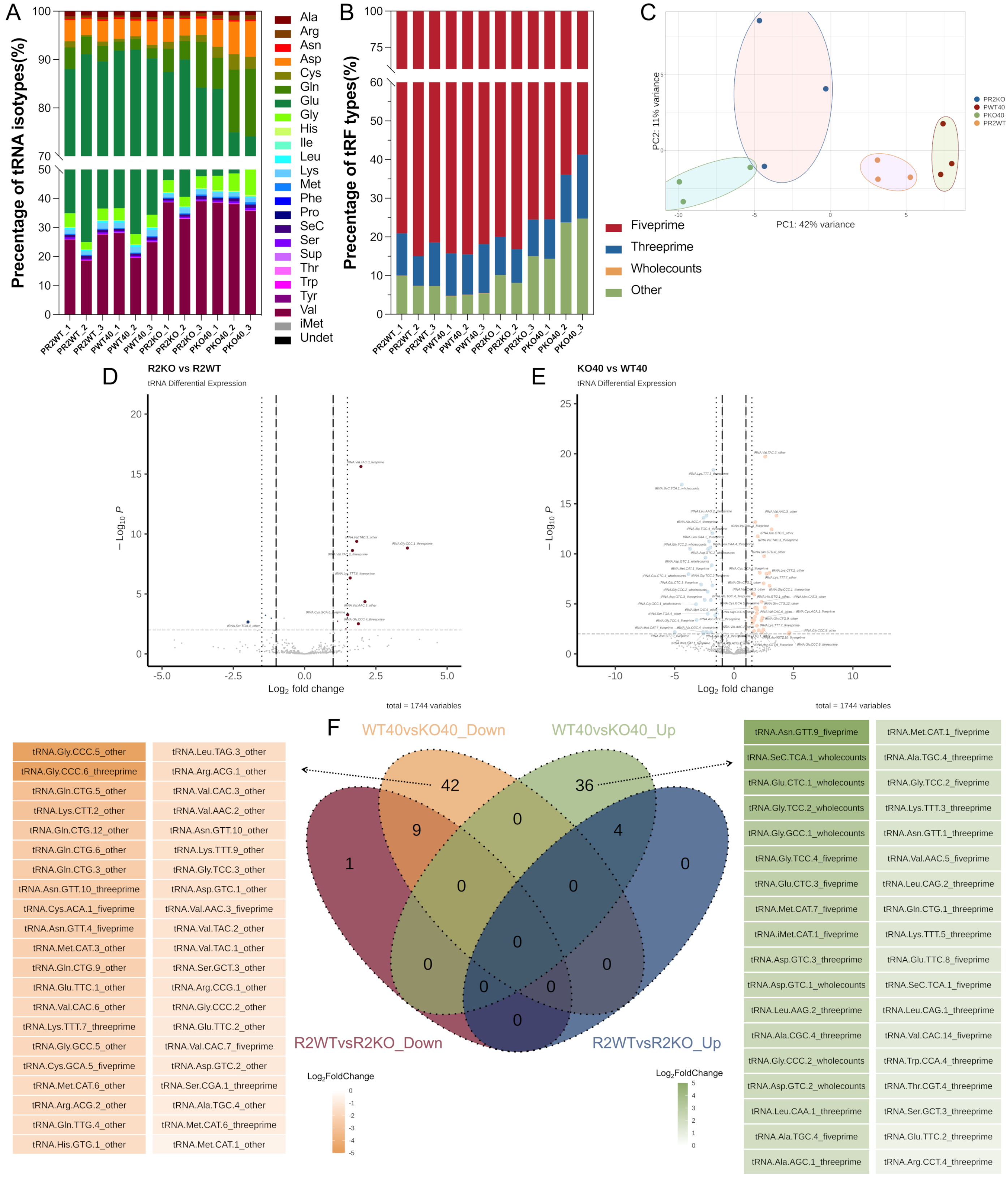


**Supplemental Figure S10. Comparison of tRNA & tDRs population obtained by 2’3’C>p amplification in Library 3**.

**(A)** Percentage of read for tRNA isotypes.

**(B)** Percentage of read for tDRs types.

**(C)** PCA plot of tDRs among 4 group.

**(D and E)** Volcano plot for transcripts with different two-by-two comparison**.** The significantly down/up-represented tRFs identified from tRAX are labelled (Cut-off: $|Log_2FC|>1.5$ and padj<0.01).

**(F)** tRFs population comparison among groups**.** Venn plot of over/under-represented tRFs for WT40vsKO40 and R2WTvsR2KO. The unique tRFs list in WT40vsKO40 group were shown by heatmap.

The five-prime tRFs correspond to reads that are within 10 bases of the start position, but do not reach the end position of the tRNA sequence and the three-prime ones to reads that are within 10 bases of the end position, but do not reach the start position of the tRNA sequence according to tRAX tutorial

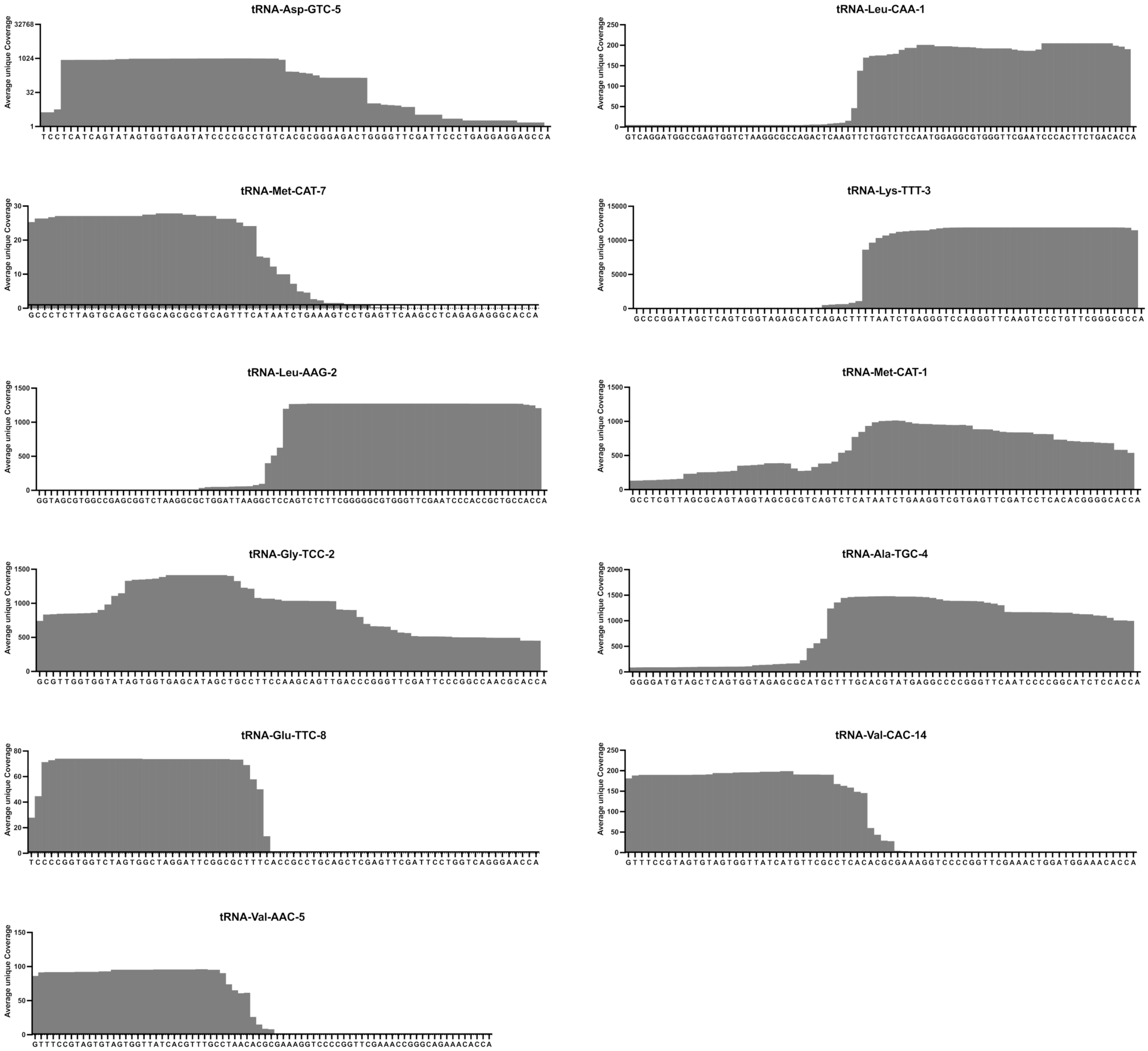


**Supplemental Figure S11: Average of unique coverage per base of selected tRNAs in WT40 group.**

The unique average coverage from WT40 group was obtained from tRAX pipeline output. According to the high to low order of log2FC in Additional file 4, the tRNAs are ordered sequentially from top to bottom, followed by left to right.

### Supplemental Table S1. tRFs in unique WT40vsKO40_up group

| | Sequences (5’-3’) | Type | tDR names* |
|---|---|---|---|
| tRNA-Ala-TGC-4 | CUUUGCACGUAUGAGGCCCCGGGUU CAAUCCCCGGCAUCUCCACC | 3’-tRFs | tDR-31:75-Ala-TGC-4 |
| tRNA-Leu-AAG-1 | CAGUCUCUUGGGGGGCGUGGGUUCG AAUCCCACCGCUGCCACC | 3’-tRFs | tDR-41:75-Leu-AAG-2 |
| tRNA-Leu-CAA-1 | UCUGGUCUCCAAUGGAGGCGUGGGU UCGAAUCCCACUUCUGACACC | 3’-tRFs | tDR-39:75-Leu-CAA-1 |
| tRNA-Lys-TTT-3 | UUAAUCUGAGGGUCCAGGGUUCAAG UCCCUGUUCGGGCGCC | 3’-halves | tDR-35:75-Lys-TTT-3 |
| tRNA-Met-CAT-7 | GCCCUCUUAGUGCAGCUGGCAGCGC GUCAGUUUC | 5’-halves | tDR-1:34-Met-CAT-7 |
| tRNA-Glu-TTC-8 | UCCCCGGUGGUCUAGUGGCUAGGAU UCGGCGCUUU | 5’-halves | tDR-1:35-Glu-TTC-3-M3 |
| tRNA-Val-AAC-5 | GUUUCCGUAGUGUAGUGGUUAUCAC GUUUGCCUAAC | 5’-halves | tDR-1:36-Val-AAC-5 |
| tRNA-Val-CAC-14 | GUUUCCGUAGUGUAGUGGUUAUCAU GUUCGCCUCAC | 5’-halves | tDR-1:36-Val-CAC-14 |

* The tDR names of the identified sequences were obtained from the tDRname (v1.3.1) pipeline
